# Cell-surface protein YwfG of *Lactococcus lactis* binds to α-1,2-linked mannose

**DOI:** 10.1101/2022.08.19.504583

**Authors:** Wataru Tsuchiya, Zui Fujimoto, Noritoshi Inagaki, Hiroyuki Nakagawa, Miwa Tanaka, Hiromi Kimoto-Nira, Toshimasa Yamazaki, Chise Suzuki

## Abstract

*Lactococcus lactis* strains are used as starter cultures in the production of fermented dairy and vegetable foods, but the species also occurs in other niches such as plant material. *Lactococcus lactis* subsp. *lactis* G50 (G50) is a plant-derived strain and potential candidate probiotics. Western blotting of cell-wall proteins using antibodies generated against whole G50 cells detected a 120-kDa protein. MALDI-TOF MS analysis identified it as YwfG, a Leu-Pro-any-Thr-Gly cell-wall-anchor-domain–containing protein. Based on a predicted domain structure, a recombinant YwfG variant covering the N-terminal half (aa 28–511) of YwfG (YwfG_28−511_) was crystallized and the crystal structure was determined. The structure consisted of an L-type lectin domain, a mucin-binding protein domain, and a mucus-binding protein repeat. Recombinant YwfG variants containing combinations of these domains (YwfG_28–270_, YwfG_28–311_, YwfG_28−511_, MubR4) were prepared and their interactions with monosaccharides were examined by isothermal titration calorimetry; the only interaction observed was between YwfG_28–270_, which contained the L-type lectin domain, and D-mannose. Among four mannobioses, α-1,2-mannobiose had the highest affinity for YwfG_28–270_ (dissociation constant = 34 μM). YwfG_28–270_ also interacted with yeast mannoproteins and yeast mannan. Soaking of the crystals of YwfG_28–511_ with mannose or α-1,2-mannobiose revealed that both sugars bound to the L-type lectin domain in a similar manner, although the presence of the mucin-binding protein domain and the mucus-binding protein repeat within the recombinant protein inhibited the interaction between the L-type lectin domain and mannose residues. Three of the YwfG variants (except MubR4) induced aggregation of yeast cells. Strain G50 also induced aggregation of yeast cells, which was abolished by deletion of *ywfG* from G50, suggesting that surface YwfG contributes to the interaction with yeast cells. These findings provide new structural and functional insights into the interaction between *L. lactis* and its ecological niche via binding of the cell-surface protein YwfG with mannose.

## Introduction

Gram-positive bacteria interact with their environment and other cells via proteins expressed on their cell surface. An important subset of these cell-surface proteins are the adhesins that mediate adhesion to other cells or environmental surfaces. For example, in the initial stages of colonization, pathogenic bacteria use their cell-surface proteins to bind to host cells [1].

In *Staphylococcus aureus*, mucus-binding proteins are anchored to the cell surface by a sortase A–mediated system that recognizes the motif LPXTG (Leu-Pro-any-Thr-Gly) [2]. In this system, the mucus-binding proteins are cleaved between the Thr and Gly of the LPXTG motif by sortase A and then undergo transpeptidation and covalent attachment to peptidoglycan in the bacterial cell wall. Sortase-anchored surface proteins fulfill key functions during the infectious process and thus sortase A is essential for host colonization and for the pathogenesis of invasive diseases. The roles of sortases and proteins containing the LPXTG motifs are well documented in pathogens, but even in nonpathogenic lactic acid bacteria a mechanism that mediates the relationship between cell surface proteins and the host that is analogous to that in pathogenic bacteria has been identified [3, 4].

Lactobacilli are Gram-positive bacteria in order *Lactobacillales*, strains of which are found in the human intestine and used in the production of fermented foods. The strains that are used as probiotics are mostly of intestinal origin and are reported to adhere to the intestinal mucus layer [5]. Understanding the interaction between lactobacilli and mucins is critical to elucidate the survival strategies of lactobacilli in a constantly changing intestinal environment, as well as to elucidate the mechanisms of their beneficial effects in the intestinal tract. Cell-surface proteins likely play a key role in these strategies and mechanisms by interacting with host receptors, such as the glycans in mucins [3, 5]. Lactobacilli express several adhesins, including moonlighting proteins (multi-functional cytoplasmic proteins) that exert adhesive functions when expressed on the cell surface [6]. Several species of lactobacilli are known to possess surface-layer (S-layer) proteins [7] that are involved in the adhesion of the cells to the host epithelium or to extracellular matrix components and in modulating the host immune response [3]. For example, knockout of S-layer protein A in *Lactobacillus acidophilus* significantly reduces binding of the mutant to DC-SIGN, a dendritic cell (DC)–specific receptor, and S-layer protein A is involved in the modulation of DC and T-cell functions [8]. Whereas certain strains of lactobacilli bind to epithelial cells via mucin in order to reside in the intestinal tract, other strains bind yeast cells used in fermentation processes [9]. In some cases, symbiosis between the yeast and lactobacilli improves fermentation [10, 11].

*Lactococcus* is another genus in order *Lactobacillales*. Strains belonging to *Lactococcus lactis* and *Lactococcus cremoris* are used as fermentation starters in the production of cheese and other dairy and vegetable fermented products. Originally, it was thought that lactococci did not reach the intestinal tract alive, but Klijn et al. [12] showed that a fraction of viable cells of a strain of *L. lactis* was able to survive passage through the human gastrointestinal tract. Recently, several lactococcal strains have been reported to have beneficial health effects such as improvement of skin status [13] and activation of human plasmacytoid DCs [14].

There are few studies on lactococcal surface proteins that interact with intestinal epithelium or DCs. *Lactococcus lactis* is used extensively in the industrial production of fermented dairy products. However, *L. lactis* strains have also been isolated from plant material [15, 16]. *Lactococcus lactis* subsp. *lactis* G50 (G50) was isolated from Napier grass and was subsequently found to have immunostimulatory activity [17]. *In vitro*, intact G50, but not heat-treated G50, can induce tumor necrosis factor α expression in the murine macrophage cell line J774.1, suggesting that a heat-sensitive molecule is involved in this induction [18]. In ovomucoid-sensitized BALB/c mice, oral administration of intact G50 cells reduces total IgE production [17]. Together, these findings indicate that G50 may have potential uses as a probiotic.

Analysis of the G50 genome sequence [19] may help reveal the cell-surface proteins that contribute to the activity of G50. Here, we detected a cell-surface protein by reaction of *L. lactis* strains with anti-G50 antibody. We used matrix-assisted laser desorption/ionization–time-of-flight mass spectrometry (MALDI-TOF MS) and X-ray crystallography to investigate the structure and interactions of this protein.

## Materials and Methods

### Strains

The lactococcal strains used in this study are listed in Table 1. The strains were grown at 30°C for 18 h in M17 (BD Difco, Sparks, MD, USA) broth containing 0.5% (wt/vol) glucose (GM17) or chemically defined medium [20] in which lactose was replaced with 2% glucose (CDMG) [21]. *Escherichia coli* BL21(DE3) (Merck, Darmstadt, Germany) was used for the expression of recombinant proteins. Three yeast strains were also used: *Saccharomyces cerevisiae* X2180-1A, *S. cerevisiae* AB9 [MAT a/α *gpi10/gpi10 ura3/URA3 leu2/LEU2*], and *S. cerevisiae* AB9-2 [MAT a/α *gpi10/gpi10 mnn2/mnn2 ura3/URA3 leu2/LEU2*] [22]. YPD containing 1% yeast extract, 2% peptone, and 2% glucose was used for culturing the yeast strains. A synthetic defined medium (SD-medium) (yeast nitrogen base without amino acids containing 0.17% ammonium sulfate [BD Difco], 2% glucose, 0.13% casamino acids [BD Difco]) was used for yeast culture for mannoprotein preparation.

**Table 1.**
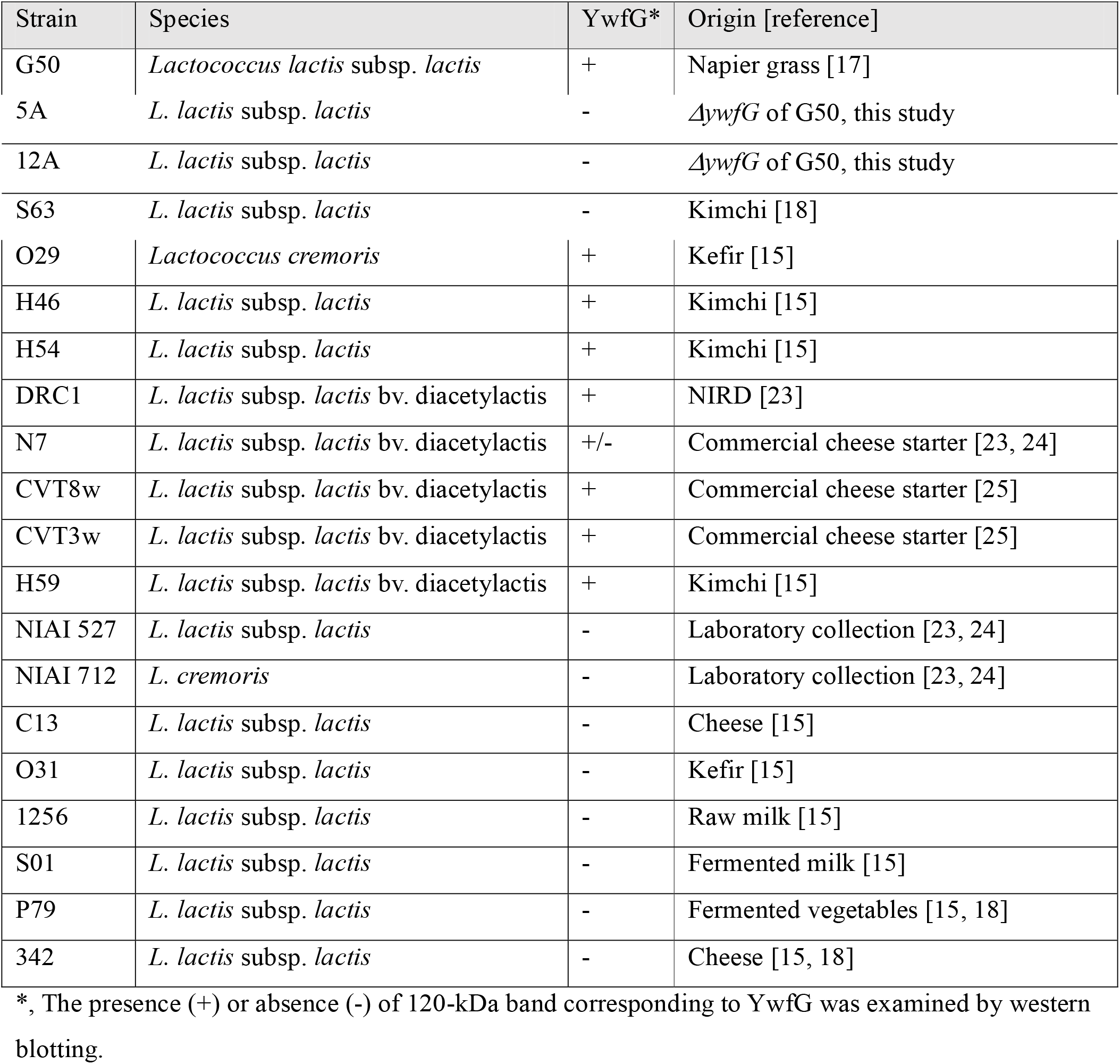
Lactococcal strains used in this study

### Preparation of cell-wall proteins, surface-exposed proteins, and cell lysates

Cell-wall proteins were extracted from the cell-wall fraction by using lysozyme, as reported by Meyrand et al. [26]. Briefly, *L. lactis* cells were cultured to an optical density (OD_600_) of 1.2 in 6 mL of GM17 medium at 30°C and centrifuged at 5,000 × *g* for 10 min at 4°C. The cell pellet was washed once with 50 mM Tris-HCl (pH 7.0) and resuspended in 1 mL of 50 mM Tris-HCl (pH 7.0) containing protease inhibitors (cOmplete, Roche Applied Science, Mannheim, Germany). The cells were disrupted with glass beads using a FastPrep system (Qbiogene, Montreal, Quebec, Canada) set at 6.0 m s^−1^ for 40 s and centrifuged at low speed (1,000 × *g*) for 2 min at 4°C to remove the glass beads and any unbroken bacteria. The supernatant was centrifuged at 16,000 × *g* for 15 min at 4°C. The pellet containing the bacterial cell envelopes was digested with lysozyme (1 mg L^−1^) and mutanolysin (200 U mL^−1^) in 50 mM Tris-HCl (pH 7.0) containing 5 mM MgCl_2_ for 3 h at 37°C, centrifuged at 10,000 × *g* for 10 min at 4°C, and the supernatants were analyzed by western blotting.

Surface-exposed proteins were prepared according to the shaving method [26]. Bacterial cells were digested by trypsin in the presence of 1.2 M sucrose and 1 mM CaCl_2_, centrifuged at 20,000 × *g* for 10 min, and supernatants were used for western blotting.

To prepare whole cell lysates, bacterial cultures (5 mL) were centrifuged, pellets were washed twice with phosphate buffer saline (PBS), and suspended in 300 μL of PBS containing 0.1% SDS, protease inhibitor cocktail (cOmplete), and Lysing matrix B (MP Biomedicals, Santa Ana, USA). Cells were disrupted with glass beads using a FastPrep system as described above, centrifuged at 13,000 × *g* for 10 min, and the supernatants were used for western blotting.

### SDS-PAGE and western blotting

SDS-PAGE was performed in a 5%–20% linear gradient gel (Atto, Tokyo, Japan). Proteins were visualized by Coomassie Brilliant Blue staining (CBB) or western blotting. After SDS-PAGE, proteins were electrotransferred to a polyvinylidene difluoride membrane (Millipore, Burlington, MA, USA) using a semi-dry transfer blotter (Atto). Rabbit polyclonal antiserum to G50 was generated against intact G50 cells and purified by using a recombinant protein A column (rProtein A Sepharose FF, Cytiva, Tokyo, Japan). The membrane was incubated with anti-G50 antiserum (1:3,000 dilution) and horseradish peroxidase–linked anti-rabbit immunoglobulin (Bio-Rad Laboratories Inc., Hercules, CA, USA; 1:5,000 dilution), and the blot was developed with an ECL Prime western blotting kit (Cytiva).

### Protein identification by MALDI-TOF MS and database searching

CBB-stained bands corresponding to the bands detected by the anti-G50 antiserum were excised from the gel and treated by in-gel digestion with a proteomics-grade trypsin (Sigma-Aldrich, St Louis, MO, USA). The digested samples were purified by using ZipTip C18 (Merck). The purified peptides were mixed with an equal volume of α-cyano-4-hydroxycinnamic acid (4-CHCA) (Shimadzu GLC Ltd., Tokyo, Japan) matrix solution (prepared as 10 mg/mL in 50% acetonitrile containing 0.1% TFA), and an aliquot of the mixture (0.5 μL) was spotted onto an ABI 4800 MALDI plate (SCIEX, Framingham, MA, USA). The mass spectrometer, a MALDI-TOF/TOF mass spectrometer (4800 Plus TOF/TOF analyzer, SCIEX) was tuned and calibrated using a calibration standards mixture (Peptide Calibration Standard II, Bruker Daltonics GmbH & Co. KG, Bremen, Germany) prior to the measurements. After the spot had been allowed to dry at room temperature, MS and MS/MS spectra were obtained. Proteins were identified using a Mascot search engine (ver. 2.4.0.; Matrix Science, London, UK) and the NCBI database.

### Protein sequence and structure analysis

Protein sequence analysis and domain fold prediction were performed on the Pfam (http://pfam.xfam.org) and the Phyre2 (http://www.sbg.bio.ic.ac.uk/phyre2)servers. Protein structures were compared on the Dali server (http://ekhidna2.biocenter.helsinki.fi/dali/)[27].

### Preparation of YwfG variants

Total bacterial DNA was isolated by using a DNeasy Blood & Tissue Kit (Qiagen, Hilden, Germany). Primers were designed using the coding sequence (CDS) of locus tag LLG50_11005 of the G50 genome sequence (accession number CP025500.1) (Table S1).

DNA fragments encoding variants of YwfG (residues 28–270, 28–336, 28–511, and 860–1034) were amplified by polymerase chain reaction using the primer sets Fw28–Rv270, Fw28–Rv336, Fw28–Rv511, and Fw860–Rv1034, respectively. The resulting fragments were inserted in NdeI/BamHI-digested pET28b (Merck) by In-Fusion cloning reactions (In-Fusion HD Cloning Kit, Takara Bio, Kusatsu, Japan) with an N-terminal His_6_-tag. The resulting constructs were transformed into *E. coli* BL21(DE3). Recombinant YwfG variants were expressed at 16°C overnight in the presence of 0.2 mM isopropyl β-D-thiogalactoside and purified by affinity chromatography using a nickel-chelating column (HisTrap HP, Cytiva), followed by gel filtration using a HiLoad 26/600 Superdex 75 pg column (Cytiva) and cation exchange chromatography using a Mono S 10/100 GL column (Cytiva). The obtained recombinant YwfG variants were named YwfG_28–270_, YwfG_28–336_, YwfG_28–511_ (numbers indicate amino acid residues), and MubR4 (residues 860–1034).

### Preparation of yeast mannoproteins

*gpi10* mutants of *S. cerevisiae* secrete mannoproteins into the culture medium [22]. Mannoproteins of AB9 strain (wild-type) and of AB9-2 strain (*mnn2* mutant lacking α-1,2-mannosyltransferase) were prepared. Yeast cells were cultured in the SD-medium for 6 days at 30°C. The cultures were centrifuged at 12,000 × *g* for 10 min at 4°C to obtain a cell-free supernatant. The mannoproteins were concentrated using a 10,000 molecular-weight-cut-off ultrafiltration unit (Centriprep, Merk Millipore, Cork, Ireland), precipitated by adding 2 volumes of ethanol, and dissolved in 20 mM Tris-HCl (pH 7.5).

### Isothermal titration calorimetry (ITC) analysis

A MicroCal iTC200 system (Malvern Panalytical, Almelo, Netherlands) was used to measure enthalpy changes associated with interactions between the YwfG variants and the following yeast cell-wall components: the monosaccharides D-glucose, D-galactose, D-mannose, D-fucose, *N*-acetyl-D-glucosamine (GlcNAc), and 2-(acetylamino)-2-deoxy-D-galactose (GalNAc) (Sigma-Aldrich); α-1,2-, α-1,3-, α-1,4-, and α-1,6-mannobiose (Dextra Laboratories, Reading, UK); mannan from *S. cerevisiae* (Sigma-Aldrich); and yeast mannoproteins.

A solution of YwfG variant (100 μM YwfG_28–270_, YwfG_28–336_, YwfG_28–511_, or MubR4 in 20 mM Tris-HCl [pH 7.5]) was placed in a 200-μL calorimeter cell, and test solution (10 mM for monosaccharides, 2 mM for monosaccharides and mannobioses, 10 mg mL^−1^ for yeast mannan, OD_280_ 1.0 for mannoproteins) was loaded into the injection syringe. The test solution was titrated into the sample cell as a sequence of 20 injections of 2-μL aliquots each. All experiments were performed at 25°C.

### Crystallization, data collection, and structure refinement

Purified YwfG_28–511_ was concentrated to 12.0 mg mL^−1^ and crystallized by the sitting-drop vapor diffusion method at 20°C using a precipitant solution consisting of 0.1 M sodium acetate (pH 5.0–5.6), 1.3 M Li_2_SO_4_, and 2% (w/v) polyethylene glycol 3350 (Hampton Research, Aliso Viejo, CA, USA). We used 50 μL of the reservoir solution and a drop consisting of 0.5 μL of the protein solution and 0.5 μL of the reservoir solution; plate-type crystals appeared within 2–3 weeks.

Diffraction data were collected at two beamlines: BL-5A housed at the Photon Factory and BL-NW12 housed at the Photon Factory Advanced Ring (both at the High Energy Accelerator Research Organization, Tsukuba, Japan). Diffraction data were collected using a Pilatus3 S6M or Pilatus3 S2M detector (Dectris, Philadelphia, PA, USA). Crystals were collected in a 0.5-mm nylon cryoloop (0.5 mm, Hampton Research), soaked in a precipitant drop containing 30% (v/v) glycerol, and cryocooled in a nitrogen gas stream to 95 K. Crystals in complex with D-mannose or α-1,2-mannobiose were obtained by adding 0.5 μL 10% D-mannose or α-1,2-mannobiose solution to the crystal drop a few minutes prior to the diffraction experiment. The data were integrated using the XDS program package [28] and scaled using the SCALA program [29] within the CCP4 program suite [30].

The ligand-free structure of YwfG_28–511_ was determined by the molecular replacement method using the Molrep program [31] with the lectin domain (residues 258–492) of SraP (PDB code 4m01 [32]) as the search model. Sugar complex structures were built using the ligand-free structure as the starting model. Manual model building and molecular refinement were performed using Coot [33] and Refmac5 [34]. Data collection and refinement statistics are shown in Table 2. The stereochemistry of the models was analyzed using the MolProbity program [35] and structural drawings were prepared using the PyMOL (https://www.pymol.org/pymol.html) or Cuemol2 program (http://cuemol.osdn.jp/en/).

**Table 2.** Data collection and refinement statistics of the YwfG fragment structures

| | Native | D-mannose complex | α-1,2-mannobiose complex |
| --- | --- | --- | --- |
| PDB code | 7YL4 | 7YL5 | 7YL6 |
| *Data collection* |  |  |  |
| Space group | *I*222 | *I*222 | *I*222 |
| Cell parameters (Å) | $a = 43.2$<br>$b = 272.9$<br>$c = 320.3$ | $a = 43.3$<br>$b = 273.2$<br>$c = 322.8$ | $a = 43.5$<br>$b = 276.6$<br>$c = 321.8$ |
| X-ray source | PF-AR BL-NW12 | PF BL-5A | PF-AR BL-NW12 |
| Wavelength (Å) | 1.00000 | 1.00000 | 1.00000 |
| Resolution (Å) | 48.6–2.50 (2.64–2.50) | 48.7–3.00 (3.16–3.00) | 43.06–2.95 (3.11–2.95) |
| No. of reflections | 871 880 (127 582) | 507 098 (73 143) | 546 645 (84 290) |
| Unique reflections | 66 894 (9 601) | 39 549 (5 675) | 42 029 (6 070) |
| Completeness (%) | 100.0 (100.0) | 100.0 (100.0) | 99.9 (100.0) |
| Multiplicity | 13.0 (13.3) | 12.8 (12.9) | 13.0 (13.9) |
| *R*-merge | 0.093 (0.810) | 0.145 (1.053) | 0.203 (1.187) |
| Average *I*/σ | 21.3 (3.4) | 16.0 (2.7) | 14.0 (2.5) |
| *Refinement* |  |  |  |
| Resolution (Å) | 45.5–2.50 (2.57–2.50) | 45.6–3.00 (3.08–3.00) | 41.5–2.95 (3.03–2.95) |
| *R*-factor | 0.246 (0.385) | 0.227 (0.365) | 0.211 (0.361) |
| *R*-free | 0.280 (0.395) | 0.296 (0.392) | 0.268 (0.387) |
| No. of water molecules | 74 | 33 | 41 |
| Average *B*-value (Å<sup>2</sup>) | 65.1 | 79.9 | 65.2 |
| r.m.s.d. from ideals |  |  |  |
| lengths (Å) | 0.011 | 0.021 | 0.014 |
| angles (°) | 1.75 | 2.28 | 2.03 |
| Ramachandran plot (%)<br>(favored/allowed/disallowed) | 94.7 / 5.0 / 0.3 | 93.7 / 5.5 / 0.8 | 95.4 / 4.3 / 0.3 |

\*Values in parentheses refer to the highest-resolution shell. r.m.s.d., root-mean-square deviation.

### Interaction of yeast cells and YwfG variants and lactococcal cells

*Saccharomyces cerevisiae* X2180-1A was cultured in YPD medium to an OD_600_ of 0.6. In microplate wells, 50 μL yeast culture, 10 μL each recombinant YwfG protein (0.5 μM or 1 μM as indicated in Fig 7), and 40 μL of PBS, D-mannose (0.5 mM), methyl α-D-mannoside (0.5 mM), or each mannobiose (1 mM) were mixed and left to settle for 1–2 h, and aggregation was observed.

Lactococcal cells were cultured in CDMG and suspended in PBS to an OD_600_ of 1.0. Then, 60 μL of each yeast and the lactococcal cells were added to 80 μL PBS and left to settle for 1–2 h, and aggregation was observed.

### Deletion of *ywfG* gene in *L. lactis* G50

The *ywfG* of *L. lactis* G50 was deleted using the pG^+^host-based allelic exchange method [36-38]. Construction of the plasmid (pCS339) used for *ywfG* deletion is described in S1 Fig. *L. lactis* G50 was treated with lithium acetate and dithiothreitol [39] and used for electroporation of pCS339. Chromosomal *ywfG* deletion *(ΔywfG*) strains were produced according to the method of Oxaran et al. [37]. Briefly, erythromycin-resistant colonies grown at 30°C were incubated in the presence of erythromycin at 37°C, which is a non-permissive temperature for pG^+^host replication in strains with pCS339 integrated into the chromosome. Cells were incubated in the absence of erythromycin at 30°C to induce plasmid excision, and then erythromycin-sensitive strains were selected at 37°C. *ywfG* deletion was confirmed by PCR and western blotting (S1 Fig).

## Results

### Identification of cell-surface proteins that reacted with anti-G50 antibody

Five lactococcal strains showing high immunostimulatory activity in our previous study (G50, S63, O29, H46, and H54) [18] were selected. Cell-wall proteins were prepared from the cell-wall fraction by using lysozyme and analyzed by western blotting by using an antibody generated against live G50 cells (Fig 1). A 37-kDa band was detected in all strains, and a strong 120-kDa band was detected in G50, O29, H46, and H54, but not in S63.

**Fig 1.**
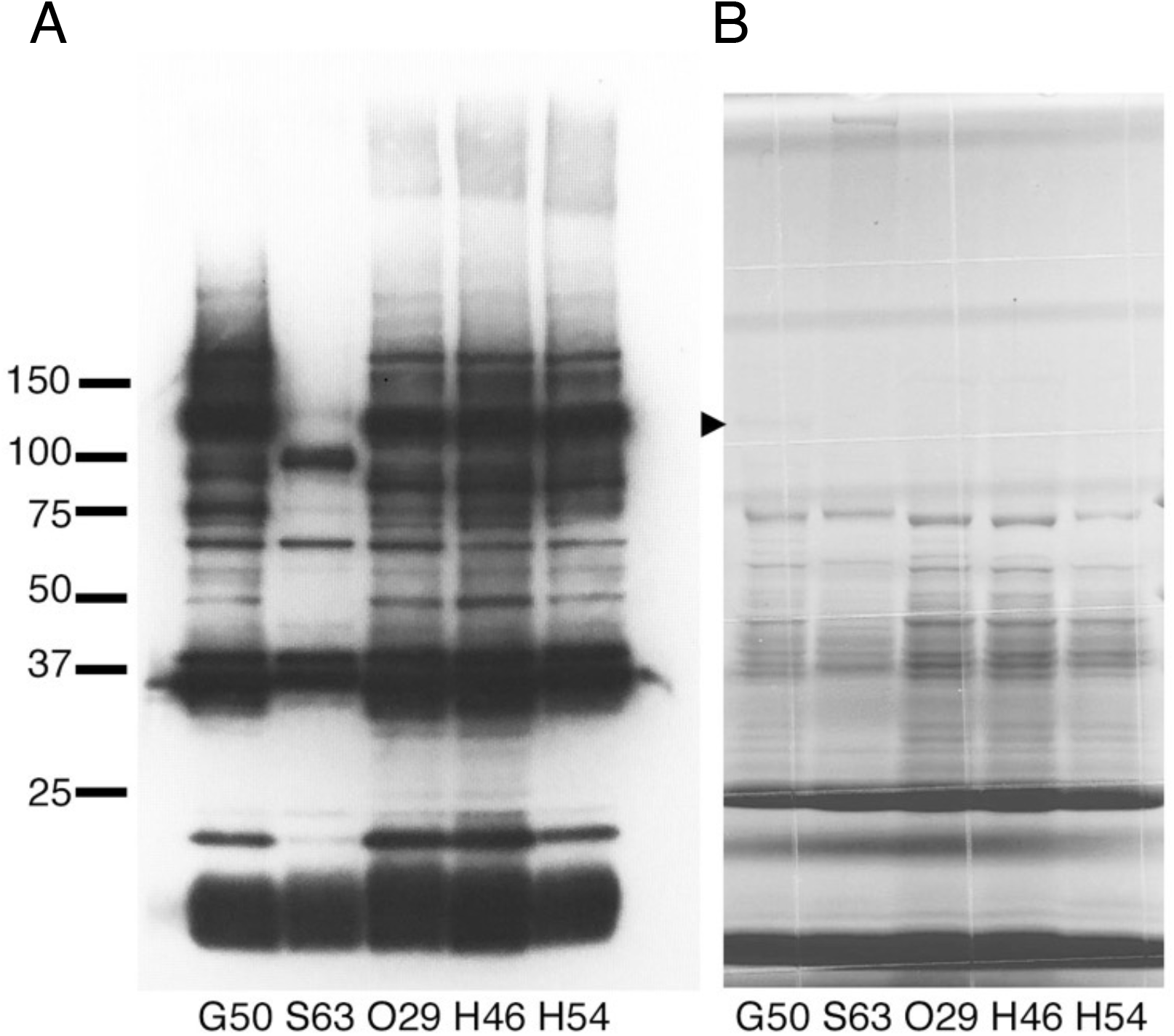
Western blot and Coomassie Brilliant Blue staining of lactococcal cell-wall proteins. (A) Western blot of cell-wall proteins of five lactococcal strains using anti-G50 antibody. (B) Coomassie Brilliant Blue staining of the same samples. The black triangle indicates the thin band of the 120-kDa protein.

In addition, a surface-exposed protein fraction was prepared from five strains (G50, S63, O29, P79, and 342) by trypsin cleavage of cell surface–exposed proteins in the presence of 1.2 M sucrose. A 120-kDa band was detected in G50 and O29 (S2A Fig). Cell lysates of 13 strains were also analyzed by western blotting (S2B Fig). The results suggested that YwfG is located both at the cell surface and in the cytoplasm, so not all of the expressed YwfG is covalently bound to the cell wall. Among the 18 lactococcal strains examined, the 120-kDa band was detected in 9 strains (Table 1).

The CBB-stained 120-kDa and 37-kDa bands from the fraction of cell-wall proteins were excised, subjected to in-gel trypsin digestion, and analyzed by MALDI-TOF MS and a Mascot search. The obtained peptide fragments from the 120-kDa protein were all aligned to the translated sequence of a CDS (LLG50_11005, GenBank: AUS70730.2) of the G50 genome (S3 Fig). This protein consists of 1107 residues and contains mucus-binding protein repeats and the LPXTG motif. The peptide fragments from the 120-kDa protein from strains O29, H46, and H54 were almost the same as those from strain G50. Interestingly, a C-terminal peptide sequence DSKPTDV<u>LPSTG</u>DSQK containing LPSTG was detected in the 120-kDa proteins of strains G50, O29, H46, and H54, suggesting that this LPXTG-motif is not used for anchoring the protein to the cell wall.

Amino acid sequence analysis using the Phyre2 and Pfam servers indicated that the 120-kDa protein contained a leguminous-type (L-type) lectin domain (Lec), a mucin-binding protein domain (Mbp), and four mucus-binding protein repeats (MubRs), followed by a C-terminal LPXTG motif (Fig 2). From its similarity to the LPXTG cell-wall-anchor-domain–containing protein of *L. lactis* subsp. *lactis* Il1403 (GenBank: AYV53882.1) [40], we named the protein YwfG, although the N-terminal 180 amino acid residues of YwfG of G50, including the lectin domain, are missing from that of Il1403.

**Fig 2.**
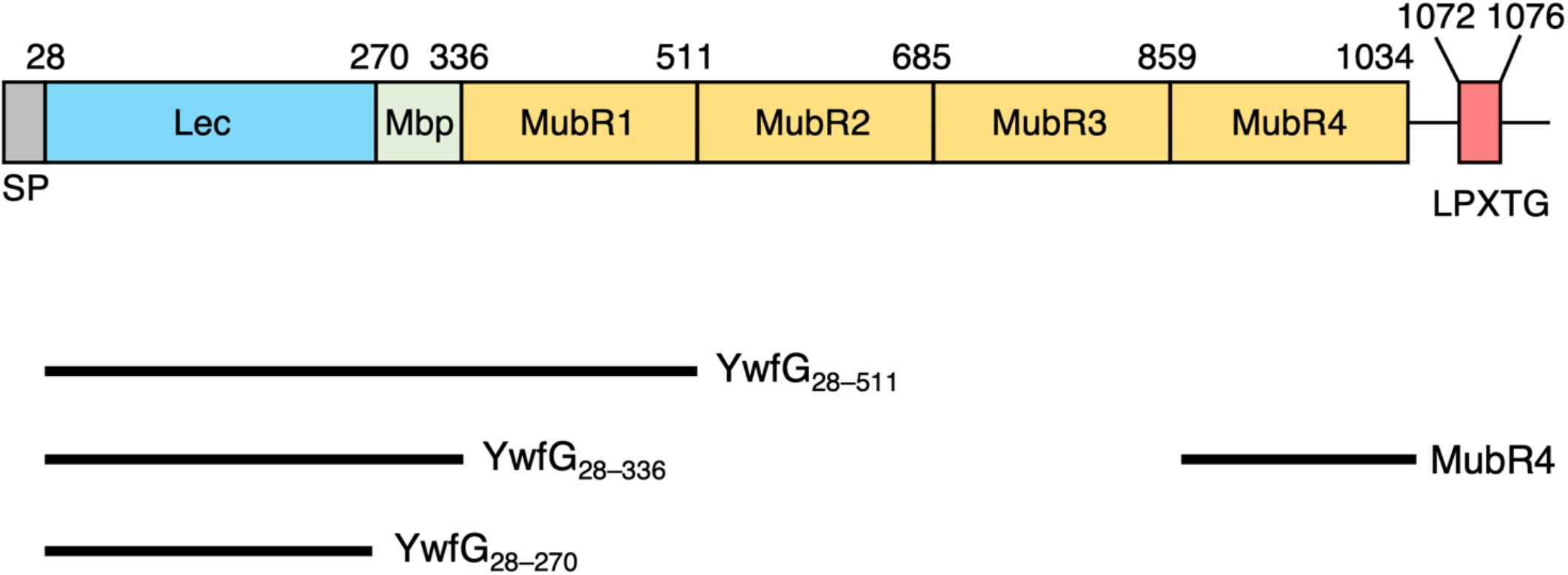
Schematic drawing of YwfG and constructions of variants. (Top) Predicted domain organization of YwfG. (Bottom) Recombinant variants constructed in the present study.

A Mascot search suggested that the 37-kDa protein was a phosphoglycerate kinase (GenBank: AUS68744.1). Because phosphoglycerate kinases have been reported to be cell-surface adhesins in *Ligilactobacillus agilis* [41] and Streptococci [42], the present lactococcal phosphoglycerate kinase could possibly function as a moonlighting protein on the cell surface.

### Interactions of the YwfG variants with monosaccharides and mannobioses

The expression of the whole YwfG protein has been attempted but expression and stability were so low that the quantity and purity of the protein could not be achieved for use in experiments. Then, four recombinant YwfG variants (YwfG_28–270_, Lec domain; YwfG_28–336_, Lec and Mbp; YwfG_28–511_, Lec, Mbp, and MubR1; and MubR4) (Fig 2) were expressed and purified. The binding of the four variants with six monosaccharides (D-glucose, D-galactose, D-mannose, D-fucose, GlcNAc, and GalNAc) and methyl α-D-mannoside was examined using ITC. The only combination of variant and monosaccharide that showed an interaction was YwfG_28–270_ with D-mannose; that is, the Lec domain alone could bind to D-mannose (S4 Fig). The binding of the three variants to four mannobioses (α-1,2-, α-1,3-, α-1,4-, and α-1,6-mannobiose) was also examined using ITC. The thermodynamic parameters for the interactions observed in the ITC assay are shown in Table 3. No binding of YwfG_28–336_, or YwfG_28–511_ to any mannobiose was observed. YwfG_28–270_ bound to all of the mannobioses, but the binding capacity of α-1,2-mannobiose was the greatest; the dissociation constant of α-1,2-mannobiose (*K*_D_ = 34 μM) was much lower than that of D-mannose (*K*_D_ = 500 μM).

**Table 3.** Thermodynamic parameters from isothermal titration calorimetry assay

| Substrate | N (sites) | $K_D$ ($\mu\text{M}$) | $\Delta H$ (cal/mol) | $\Delta S$ (cal/mol/deg) |
| --- | --- | --- | --- | --- |
| D-mannose* | (1.00) | $500 \pm 37$ | $-2098 \pm 83.59$ | 8.07 |
| α-1,2-mannobiose | $1.19 \pm 0.0104$ | $34.4 \pm 0.97$ | $-5236 \pm 62.07$ | 2.86 |
| α-1,3-mannobiose | $0.757 \pm 0.148$ | $172 \pm 21$ | $-6073 \pm 1387$ | -3.15 |
| α-1,4-mannobiose | $1.82 \pm 0.171$ | $328 \pm 91$ | $-2206 \pm 309.7$ | 9.55 |
| α-1,6-mannobiose | $1.21 \pm 0.0839$ | $210 \pm 13$ | $-4235 \pm 372.8$ | 2.62 |

\* Thermodynamic parameters determined with fixed stoichiometry ( $N = 1$ ).
N, stoichiometry; $K_D$ , dissociation constant; $\Delta H$ , enthalpy change; $\Delta S$ , entropy change.

### Crystal structure of YwfG_28–511_

Next, we elucidated the crystal structure of recombinant YwfG_28–511_. We solved it by the molecular replacement method at 2.50-Å resolution (Table 2). The crystal belonged to space group *I*222 with two molecules (I and II) per asymmetric unit. Molecule I was modeled with continuous residues from 40–511, whereas only residues 41–417 and 446–465 were modeled in molecule II. The electron density map of N-terminal residues 28–39 could not be modeled in either molecule. YwfG_28–511_ fragment had an overall elongated shape with an overall diameter of 40 Å and length of 180 Å, comprised 35 β-sheets and 4 α-helices, and was formed by head-to-tail tandem connection of the three domains, Lec, Mbp, and MubR1 (Fig 3A, B). Compared with molecule I, molecule II had a more bent structure with a larger angle between domains (Fig 3C).

**Fig 3.**
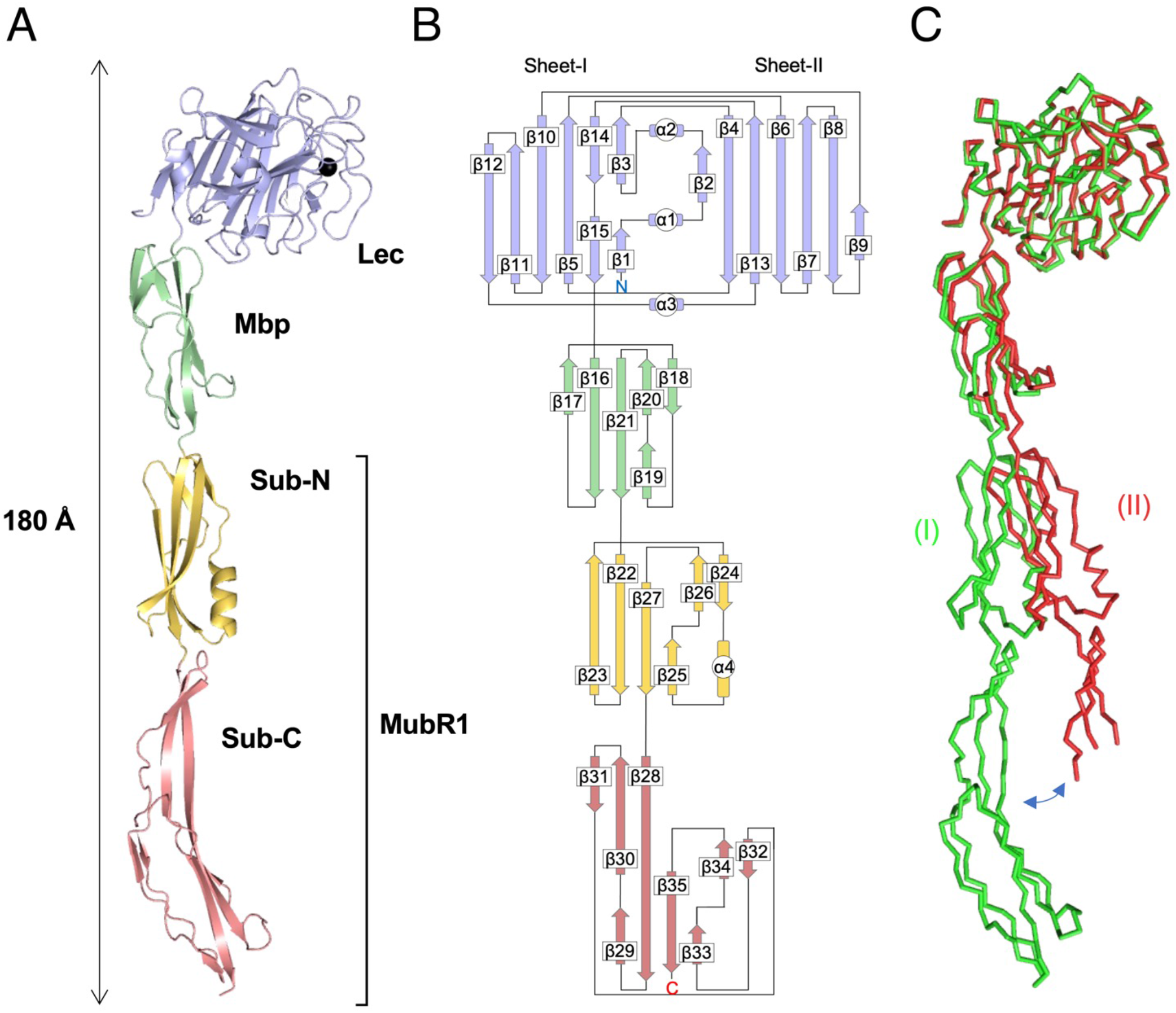
Overall structure of YwfG_28–511_. (A) X-ray crystal structure and (B) topology diagram of YwfG_28–511_ molecule I. The Lec, Mbp, Sub-N, Sub-C domains, and calcium ion are colored blue, green, orange, red, and black, respectively. (C) Superimposition of the crystal structures of YwfG_28–511_ molecules I (green) and II (red).

Starting from the N-terminus, the first domain (residues 41–268) had a jelly roll–type β-sandwich structure comprising 15 β-strands and 3 short α-helices. The core structure consisted of an antiparallel β-sheet fold formed from two sheets: sheet-I (β1, β3, β5, β10-12, β14, and β15) and sheet-II (β2, β4, β6-β9, and β13). A calcium-binding site was observed between sheet-II and the loop connecting the β7 and β8 strands (loop 7–8). One Ca^2+^ ion was coordinated by Asp152 (Oδ1 and Oδ2) belonging to β7; Tyr154 (O), Asn156 (Oδ1), and Asp166 (Oδ1) located in loop 7–8; and one water molecule. Structural similarity analysis using the Dali server [27] revealed that this domain had the highest similarity to *S. aureus* SraP lectin domain (PDB code 4m01), with a structural alignment Z-score of 35.5 and root-mean-square deviation (r.m.s.d). of 1.4 Å over 226 aligned residues (Fig 4A).

**Fig 4.**
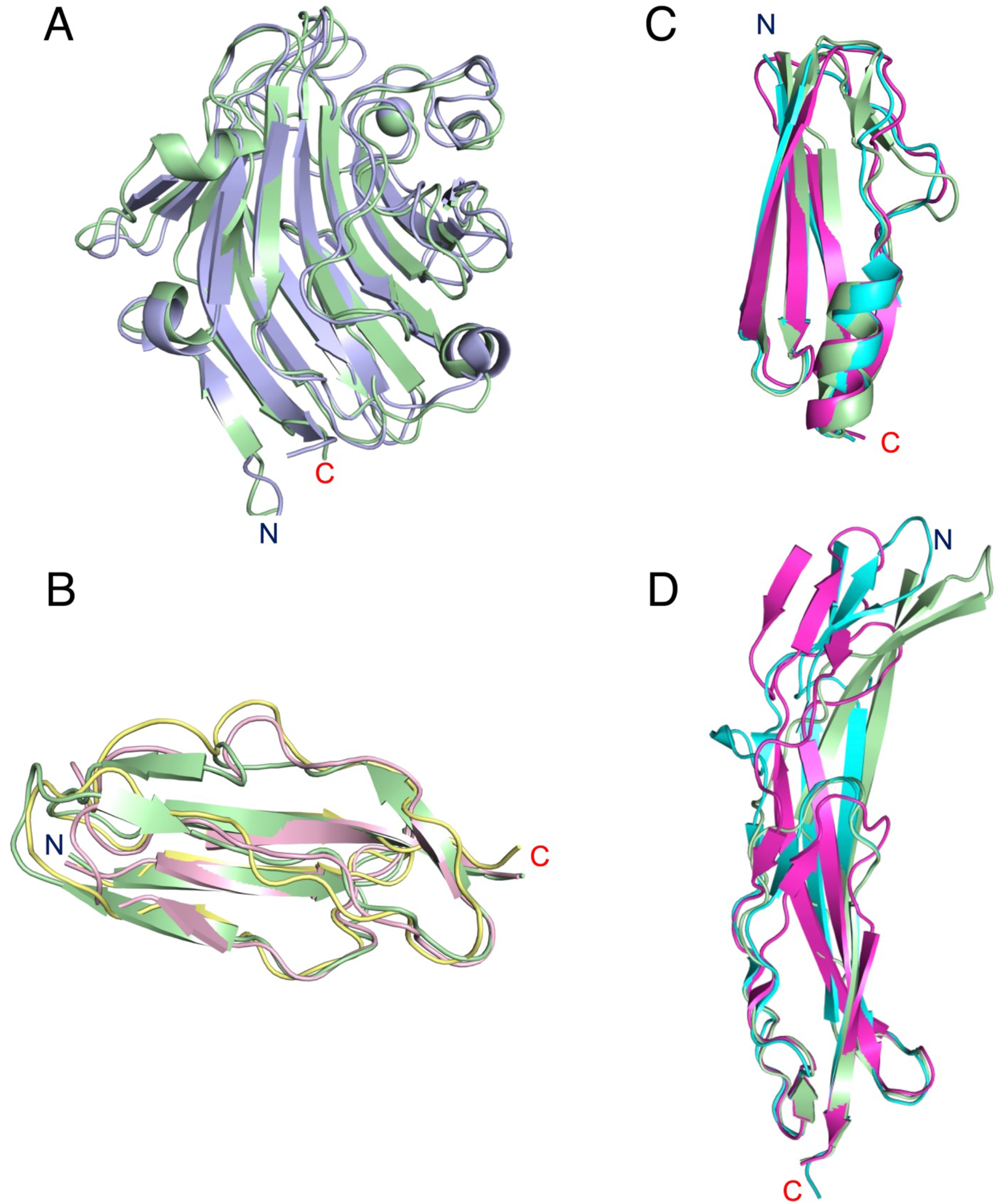
Comparison of YwfG domain structures. (A) Superimposed ribbon models of the Lec domain of YwfG (green) and *Staphylococcus aureus* SraP (light blue, PDB code 4m01). (B) Superimposition of the Mbp domain structures of YwfG (green) and *L. monocytogenes* Imo0835 (yellow, PDB code 2kvz; pink, PDB code 2kt7). Superimposition of the (C) Sub-N and (D) Sub-C structures of YwfG MubR1 on the B1 and B2 subdomains, respectively, of *L. reuteri* MubRV (cyan, PDB code 4mt5) and MubR5 (magenta, PDB code 3i57).

The second domain (residues 271–336) comprised six β-strands folded in a β-sandwich composed of a mixed β-sheet formed from β16, β17, β19, and β21 and an antiparallel β-sheet consisting of β18 and β20. This structure is similar to the Mbp domains of *Listeria monocytogenes* mucin-binding and putative peptidoglycan-binding protein Imo0835 (PDB codes 2kt7 and 2kvz, respectively) with a structural alignment Z-score of 7.7–8.2 and r.m.s.d. of 1.6–1.9 Å over 64–66 Cα atoms (Fig 4B).

The third domain (residues 337–511) had a structure in which two subdomains were connected by a three-stranded β-sheet. The N-terminal subdomain (Sub-N, residues 337–409) had a structure called a ubiquitin-like β-grasp fold, similar to that found in proteins in the immunoglobulin-binding superfamily, formed from six β-strands (β22–β27) and one α-helix (α4). The C-terminal subdomain (Sub-C, residues 410–511) consisted of a second domain-like fold formed from six β-strands (β28, β29, β32–β35) and a twisted antiparallel β-sheet (β28, β30, β31) located between the subdomains. The modeling of Sub-C in molecule (II) was incomplete because the electron density map was unclear. Structural comparison analysis revealed that the third YwfG domain showed high similarity with MubR folds of a mucus-binding protein from *Limosilactobacillus reuteri*, Mub-RV (PDB code 4mt5) [4] and Mub-R5 (PDB code 3i57) [43], with a Z-score of 12.3–14.5 and r.m.s.d. of 3.3–6.3 Å over 155–165 aligned residues. The Sub-N and Sub-C domains of YwfG were very similar to the B1 and B2 domains of the *L. reuteri* MubRs, respectively, although the β-sheet of the third domain of YwfG is more bent (Fig 4C, D).

### Substrate recognition mechanism

A structural analysis of the substrate-binding complex was performed to elucidate the substrate recognition mechanism of YwfG for the glycoconjugates with which an interaction was confirmed by the ITC assay. To obtain crystals of substrate-bound YwfG, ligand-free crystals were soaked in a solution containing D-mannose or α-1,2-mannobiose. The structures of the D-mannose– and α-1,2-mannobiose–YwfG_28–511_ complexes were determined at 3.00 Å and 2.95 Å, respectively. In each complex, a single substrate molecule was bound to the recognition site, which was located in the groove-like region between loop 7–8 and loop 13–14 on the YwfG_28–270_ surface and partially overlapped with the calcium-binding site.

In the Man–YwfG_28–511_ complex, the recognition of D-mannose by the Lec domain was the result of a hydrogen bond network involving six amino acid residues (Fig 5A). The D-mannose-OH6 group was recognized by the main-chain N of Gly250 and Phe252 and the main-chain O of Phe252 in loop 13–14. The main-chain amide of Ala251 also formed a hydrogen bond with D-mannose-O5. The D-mannose-OH4 group interacted with the sidechain of Asn156 (Nδ2) coordinating with the calcium ion and sidechain of Asp117 (Oδ1) in the β6 strand. Furthermore, the D-mannose-OH3 group interacted indirectly with the main-chain N of Gly136 in loop 6–7 via a water molecule. In addition, the aromatic sidechain of Tyr154 formed a CH-π stacking interaction with the hydrophobic face of D-mannose.

**Fig 5.**
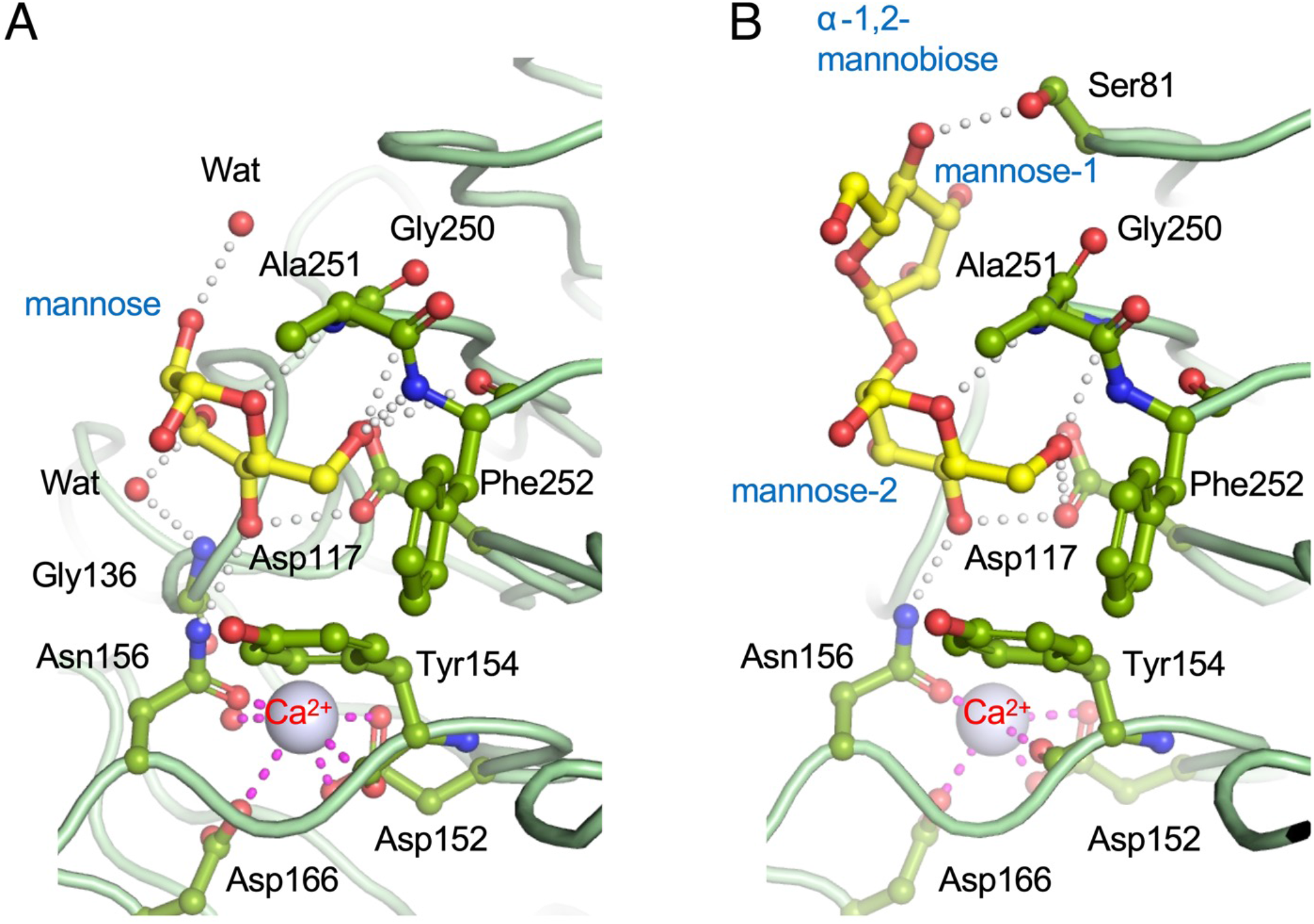
Substrate recognition by the YwfG Lec domain. (A) D-mannose and (B) α-1,2-mannobiose binding mechanism. The sugar substrate molecules and the YwfG residues that interact with these are indicated by the yellow and green ball-and-stick models, respectively. Calcium ions are shown as blue-white spheres.

The crystal structure of the α-1,2-mannobiose–YwfG_28–511_ complex showed a wider range of substrate interactions than was seen for the recognition of mannose (Fig 5B). The OH4 group of the non-reducing terminal mannose residue (mannose-1) formed a hydrogen bond with the sidechain of the Ser81 (Oγ) in loop 3–4. The reducing terminal mannose residue (mannose-2) was recognized in a manner similar to that of the monosaccharide mannose. The mannose-2-OH1 group was not involved in the recognition mechanism and was oriented toward the open space outside the ligand-binding pocket.

### Interaction of YwfG variants with yeast mannoproteins

Yeast *gpi10* mutants secrete mannoproteins into the culture media [22]. Here, secreted mannoproteins were concentrated from the culture filtrate by ultrafiltration and ethanol precipitation. Interaction of YwfG_28–270_ with mannoproteins was examined using ITC. Thermal energetic changes, which indicate an interaction, were observed for mannoproteins obtained from AB9 and AB9-2 and for the commercial yeast mannan. Yeast mannoproteins are highly glycosylated cell-wall proteins consisting of 20% protein and 80% D-mannose.

Mannoprotein from AB9 is wild-type high-mannose glycan, whereas that from AB9-2 is high-mannose glycan lacking α-1,2-linked mannose. Consistent with the observed binding capacity to α-1,2-mannobiose (Table 2), the interactions of YwfG_28–270_ with mannoprotein from AB9 and yeast mannan were more intense than that with the mannoprotein from AB9-2 (Fig 6).

**Fig 6.**
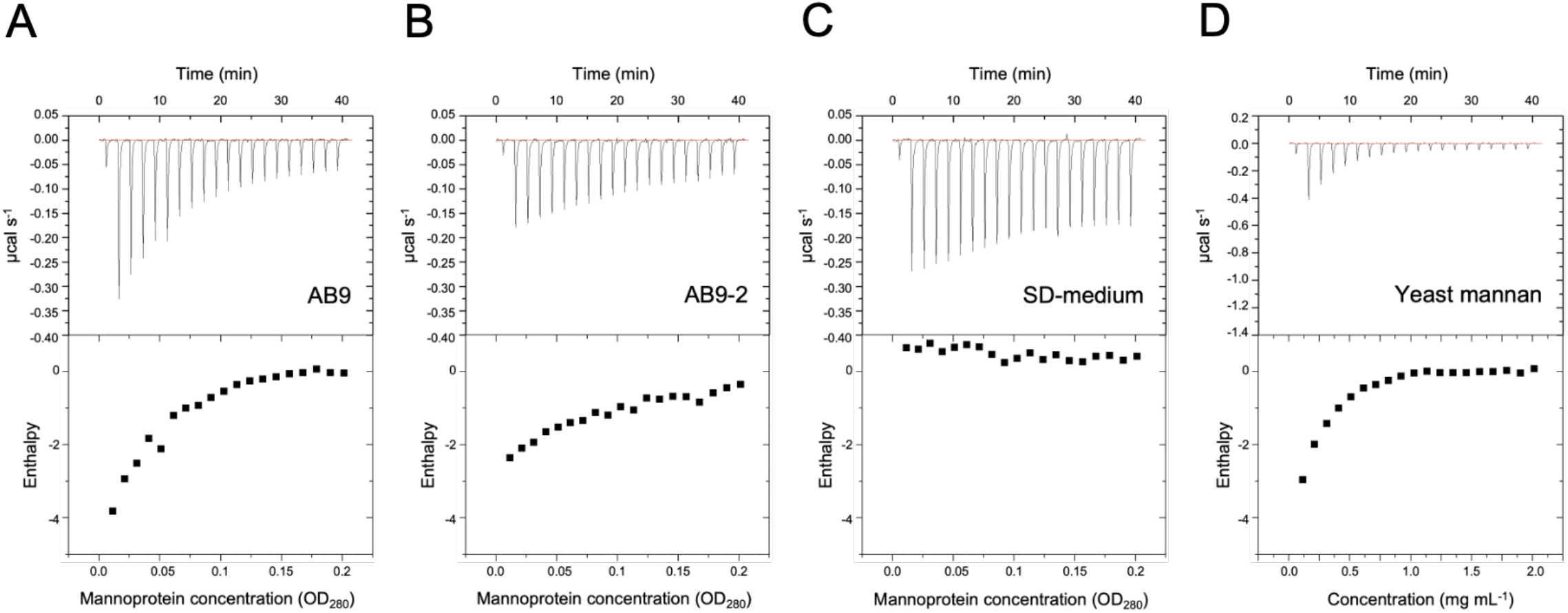
Interaction of YwfG_28–270_ with yeast mannan proteins. Isothermal titration of YwfG_28–270_ against (A) AB9 (wild type) mannoprotein, (B) AB9-2 (*mnm2* mutant) mannoprotein, (C) SD-medium, and (D) yeast mannan.

### Interaction of YwfG variants with yeast cells

The interactions of the four recombinant variants (YwfG_28–270_, YwfG_28–336_, YwfG_28–511_, and MubR4) with yeast X2180-1A, which expresses high-mannose glycans on its cell surface, were investigated (Fig 7). When X2180-1A and each recombinant variant (1 μM) were added to a well of a round-bottomed titer plate, the cells were distributed uniformly at the bottom of the wells in the presence of YwfG_28–270_, YwfG_28–336_, YwfG_28–511_, but not in the presence of MubR4 (Fig 7A). Non-aggregated yeast cells settle toward the center of round-bottomed vessels and wells, but when aggregated, they assume a uniform distribution across the bottom of the vessel or well. Such a uniform distribution or “creeping” phenotype [44] is observed in glycosylation-deficient mutants [45] and in mating yeast cells. Arras et al. [44] have described that yeast cells ‘creep’ up the sides of the culture vessel when cells of opposite mating types are mixed, and they have shown that this ‘creeping’ phenotype is a consequence of cell aggregation. The aggregation induced by YwfG_28–336_ but not by YwfG_28–270_ or YwfG_28–511_ was inhibited by the addition of 1 mM D-mannose or α methyl-D-mannoside (Fig 7B). Similarly, when 1 mM mannobiose was added, the aggregation induced by YwfG_28–336_ and YwfG_28–511_ but not by YwfG_28–270_ was clearly inhibited in the presence of α-1,2-mannobiose and was weakly inhibited in the presence of α-1,3-, α-1,4-, or α-1,6-mannobiose (Fig 7C). Aggregation was observed in the presence of *L. lactis* G50, but this aggregation was inhibited by the addition of 1 mM methyl α-D-mannoside. Furthermore, aggregation was not observed in the presence of *ΔywfG* strains (Fig 7D), indicating that cell-surface YwfG contributes to the interaction with yeast cells.

**Fig 7.**
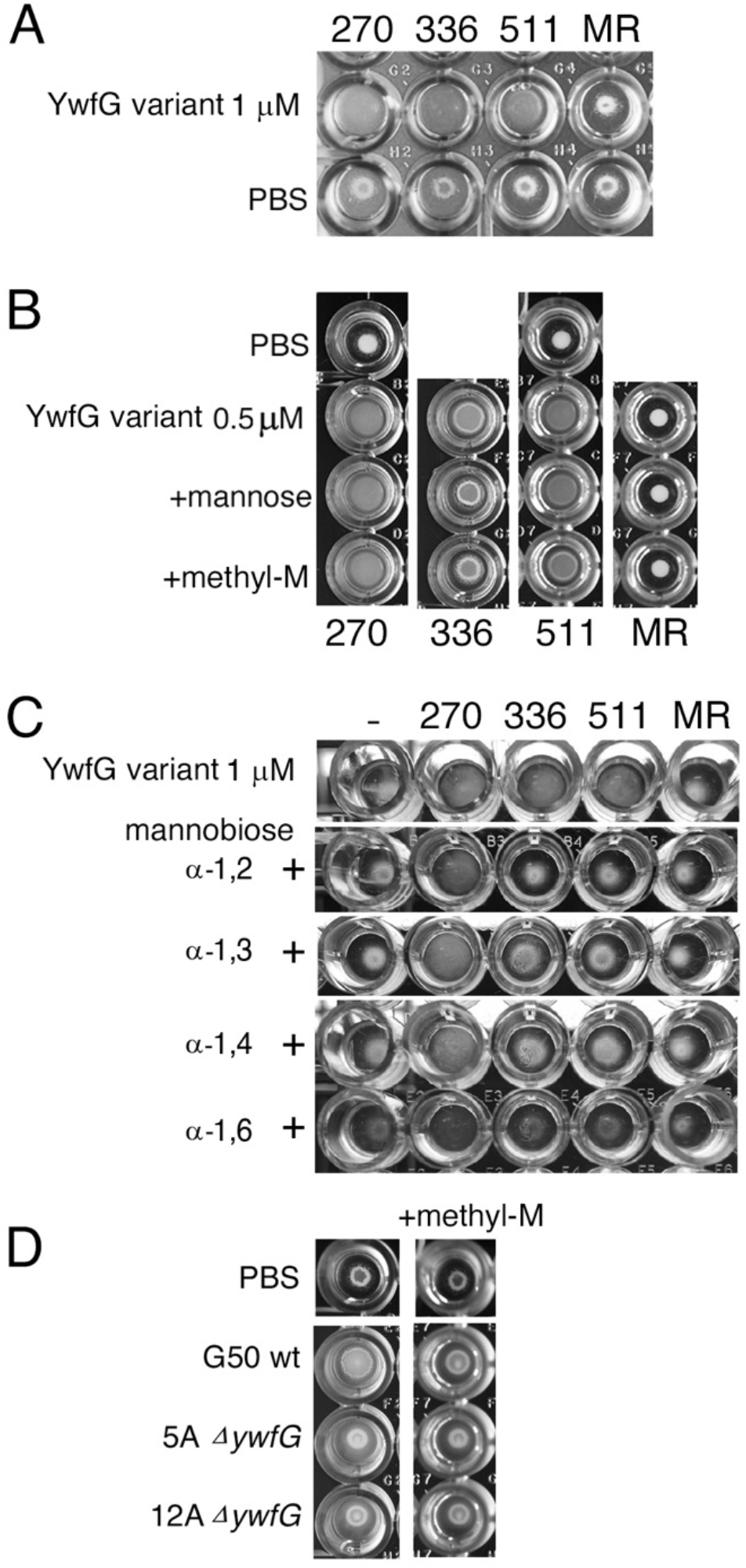
Interaction of yeast cells with YwfG variants and *L. lactis*. (A) Yeast cells in the presence of 1 μM of each variant (270 [YwfG_28–270_], 366 [YwfG_28–366_], 511 [YwfG_28–511_], MR [MubR4]) or PBS. (B) Yeast cells in the presence of 0.5 μM of each variant and 0.5 mM D-mannose or 0.5 mM methyl α-D-mannoside (methyl-M). (C) Yeast cells in the presence of 1 μM of each variant and 1 mM of each mannobiose (α-1,2, α-1,3, α-1,4, or α-1,6-mannobiose). (D) Yeast cells in the presence of *L. lactis* G50 or the *ywfG* deletion strains (5A, 12A) in the presence or absence of methyl-M.

## Discussion

Here, we found that YwfG is a cell-surface protein that reacts with a strain-specific antibody against *L. lactis* G50. This protein is reportedly to be an SrtA substrate [40]. Sortase cleaves between the Thr and Gly of this motif and covalently anchors the substrate protein to the cell wall. We detected a peptide sequence including LPSTG (DSKPTDV<u>LPSTG</u>DSQK) by MALDI-TOF MS not only in YwfG of strain G50 but also in that of the other strains examined. Since YwfG was detected in the surface-exposed protein fraction, in cell lysates, and in the cell-wall protein fraction, it is unclear whether YwfG with an uncleaved LPXTG motif is located not only in the cytoplasm but also at the cell surface. The *ΔywfG* strains did not cause yeast aggregation, suggesting that YwfG on the cell surface contributes to the aggregation or interaction with mannose residues.

Structural analysis revealed that YwfG_28–511_ is composed of three different domains: Lec, Mbp, and MubR. These domains have structural features of many gram-positive bacterial cell wall adhesins, suggesting that YwfG has a similar function [4, 43, 46, 47]. Adhesins are considered to function most effectively when their substrate-binding sites protrude as far as possible from the bacterial surface [4]. *Lactococcus lactis* G50 YwfG has a Lec domain at the tip of an approximately 600–650-Å long column consisting of one Mbp and four MubRs extending from the cell wall, suggesting that the structure is suitable for binding to distant target glycans.

The Lec domains of YwfG and *S. aureus* SraP have similar structures, although their substrate specificities differ markedly. The legume (L)-type lectin–like module of SraP specifically binds to *N*-acetylneuraminic acid and may mediate adhesion of *S. aureus* to host sialylated receptors and interaction of *S. aureus* with host epithelial cells [32]. In contrast, the ITC analysis in the present study revealed that YwfG_28–270_ exhibits high affinity for D-mannose and mannobiose, especially α-1,2-mannobiose. Crystallographic analysis of the YwfG–substrate complexes provided structural evidence of this substrate specificity. In the recognition mechanism of α-1,2-mannobiose by YwfG, mannose-2 (the reducing terminal mannose residue) was involved in multiple interactions, whereas only one hydrogen bond was observed for mannose-1 (the non-reducing terminal mannose residue), indicating that mannose-2 and mannose-1 are the major and auxiliary recognition sites, respectively.

Furthermore, the mannose-2-OH1 group is not involved in the interaction and is oriented toward the outside of the substrate-binding pocket. Since the OH1 group contributes to the polymerization of sugars, we hypothesize that the Lec domain of YwfG will also bind to longer-chain sugars possessing the mannose-α-1,2-mannose structure. This difference in the sugar-binding affinities of the Lec domains of lactococcal YwfG and the L-type lectin-like modules of staphylococcal SraP may explain the different host specificities, with the former interacting with plant substrates or yeast cells and the latter interacting with intestinal epithelium.

Because YwfG was recognized by an anti-G50 antibody that was raised against intact G50 cells, YwfG was recognized as an epitope and could interact with host immune cells. *L. reuteri* mucus-binding protein repeats, Mub-R5 and Mub-RV, which have high structural similarity to the MubR domain of YwfG, have been reported to bind not only to mucin glycans but also to mammalian secretory Ig such as IgA, IgM, and IgG at the mucosal surface of the digestive tract [43]. These interactions of the *L. reuteri* MubRs are considered to contribute to the promotion of bacterial retention in the host mucosal epithelium and the modulation of host immune responses [43]; MubR of YwfG may have similar functions. Mannose-specific adhesin (Msa) of *Lactiplantibacillus plantarum* WCSF1 also contains a MubR and an L-type lectin domain, as does YwfG, and its mannose-specific binding function is considered an essential factor for the probiotic properties of *L. plantarum* WCSF1 [46]. The lectin domains of Msa and YwfG, which contribute to mannose-specific binding, have only 37% amino acid identity. Mannose binding may be necessary for *L. lactis* to bind to plant substrates or yeast cells. Similarly, it may play an important role in interactions with mannose-containing receptors and other factors in the gastrointestinal tract. Further studies are required to elucidate the role of *L. lactis* YwfG in the host–probiotic interaction and the host response when bacterial adhesins adhere to intestinal epithelial cells or immune cells.

A Blastp search [48] showed that at least 55 lactococcal strains have YwfG homologs with more that 90% identity and query cover. YwfG is composed of a complete or incomplete N-terminal Lec domain and various numbers of MubRs. The plant-derived *L. lactis* KF147 has a CDS of YwfG (GenBank: ADA65975.1) with complete Lec, Mbp, and two MubR domains. *Lactococcus cremoris* MG1363, a plasmid-free progeny of the dairy starter strain NCDO712, has a CDS of YwfG (GenBank: CAL99029.1) with complete Lec, Mbp, and three MubR domains. The complete set of amino acid residues involved in mannose and calcium binding (D117, D152, D166, N156, Y154, and A251) are conserved in the Lec domains of YwfG of strains KF147, MG1363, and G50. YwfG was not detected in the surface proteome of *L. lactis* NZ9000, a derivative of MG1363 [49]. The authors digested cell-surface proteins with trypsin in the presence of sucrose and DTT and analyzed the resulting peptides by LC/MS, but because YwfG has no cysteine residue, it may be trypsin resistant even in the presence of DTT. Indeed, we did observe the 120 kDa band in western blots of cell surface–exposed proteins prepared by using trypsin (S2A Fig).

In contrast, *L. lactis* Il1403 has a CDS of YwfG (GenBank: AYV53882.1) lacking half (120 amino acid residues) of the N-terminal Lec domain of YwfG. The preceding CDS (GenBank: AYV53883.1) encodes 157 amino acids that are similar to the N-terminal 140 amino acid residues of YwfG of G50. There is a single-base deletion between 2259225 and 2259227 that fragments the complete YwfG sequence. It is unclear whether this is a sequencing error or an actual mutation. The YwfG of Il1403 has been confirmed to be a cell-wall-associated protein that is present in the wild-type strain and absent in the sortase A mutant [40], suggesting that those proteins are likely substrates of sortase A.

Strains of *L. lactis* are widely used in the production of fermented vegetables and various fermented dairy products, and these strains are frequently isolated from non-dairy niches, including soil and plant substrates [50]. The plant-derived *L. lactis* KF147 has gene sets for degrading complex plant polymers and is adapted to grow on substrates derived from plant cell walls [51]. Some plant cell walls contain galactomannans and glucomannans, which are polymers consisting primarily of mannose (α-D-mannosides). Degradation of mannans to D-mannose can be followed by conversion to D-fructose by mannose isomerase. In the present study, a 35-kb region (1462553 to 1497747) of the G50 genome showed more than 98% identity with 100% coverage with more than 40 *Lactococcus* strains including KF147 and Il1403. This region encodes various proteins involved in glycan degradation and transport, such as fructokinase, β-glucosidase, endo-β-*N*-acetylglucosaminidase, glucohydrolase, lacto-*N*-biosidase, sugar transporters, α-mannosidases, and xylose operon. These proteins therefore may contribute to the degradation of mannans and/or plant substrates and uptake of monosaccharides for the growth of the strains. Anchoring lectins that bind to mannose-containing glycans distributed in plant cell walls to the cell-surface layer may be a necessary strategy for degrading plant substrates.

Here, we experimentally clarified the structure and function of the lactococcal cell surface protein YwfG. Categorized as a mucus-binding protein, this protein is homologous to the *Staphylococcus aureus* SraP lectin, but unlike the lectins of the pathogenic bacteria, YwfG bound mannose. These findings provide new structural and functional insights into the interaction between *L. lactis* and its ecological niche, showing that these interactions are distinct from the bacteria–host interactions of pathogenic bacteria.

## Acknowledgements

The authors thank Ms. Tomoko Sato for her outstanding technical support in the MALDI-TOF MS analysis and Dr. E. Maguin for supplying the plasmid pG^+^host9.

## Supporting information

**S1 Table** Plasmids and primers used in this study.

**S1 Fig A** Map of the pCS339 plasmid for *ywfG* deletion. To construct this plasmid, a 984-bp upstream fragment (2240620–2241603) and a 991-bp downstream fragment (2236506–2237496) of the *ywfG* gene (LLG50_11005, complementary 2237254–2240577) were amplified by PCR; each fragment was ligated into the pGEM-T Easy plasmid, resulting in pCS336 and pCS337, respectively. The PstI fragment (967 bp) of pCS336 was ligated into the PstI site of pCS337, resulting in pCS338. SacII–digested pCS338 and pG+host9 were ligated, resulting in pCS339. **B**, Western blots of the cell-wall proteins of *L. lactis* G50, O29, H46, and *ΔywfG* strains.

**S2 Fig A** Western blots of surface-exposed proteins of lactococcal strains S63, G50, P79, 342, and O29. **B**, Western blots of cell lysates of 13 lactococcal strains.

**S3 Fig** Deduced amino acid sequence of LLG50_11005 and peptides obtained by MALDI-TOF MS analysis. Peptides obtained were aligned to the deduced sequence and are shown underlined and in bold. Peptides that were detected on both sides of the trypsin cleavage site are indicated by a triangle.

**S4 Fig Interaction of YwfG**_**28–270**_ **with monosaccharides**. Isothermal titration of YwfG_28–270_ against D-glucose, D-galactose, D-mannose, D-fucose, GlcNAc, GalNAc.

**S1_raw_Fig 1 image**. Two gels loaded with the same samples, one for CBB staining and one for western blotting, were simultaneously subjected to SDS-PAGE. Proteins were visualized by CBB staining (Left panel, Fig 1B) or western blotting visualizing by exposure with ECL regent (Right panel, Fig 1A). The middle panel is the PVDF membrane after exposure with ECL, visualizing the protein on which the ECL reagent bound.

**S2_raw_Fig 7 image**. Interaction of yeast cells and YwfG derivatives was observed in a microtiter plate. Photographing yeast cells accumulated at the bottom of the wells requires oblique lighting, and it is difficult to take a photograph all wells at once, so the photographs were staggered and cut and pasted.

**7YL4_D_1300030883_val-report-full_P1**. A PDB Summary Validation Report for 7YL4, the ligand-free structure of YwfG_28–511_.

**7YL5_D_1300030884_val-report-full_P1**. A PDB Summary Validation Report for 7YL5, the D-mannose complex structure of YwfG_28–511_.

**7YL6_D_1300030939_val-report-full_P1**. A PDB Summary Validation Report for 7YL6, the α-1,2-mannobiose complex structure of YwfG_28–511_.

## References

1. Chan JM, Gori A, Nobbs AH, Heyderman RS. Streptococcal serine-rich repeat proteins in colonization and disease. Front Microbiol. 2020;11:593356. doi: 10.3389/fmicb.2020.593356.

2. Schneewind O, Missiakas D. Sortases, surface proteins, and their roles in Staphylococcus aureus disease and vaccine development. Microbiol Spectr. 2019;7(1). doi: 10.1128/microbiolspec.PSIB-0004-2018.

3. Muscariello L, De Siena B, Marasco R. Lactobacillus cell surface proteins involved in interaction with mucus and extracellular matrix components. Curr Microbiol. 2020;77(12):3831–3841. doi: 10.1007/s00284-020-02243-5.

4. Etzold S, Kober OI, Mackenzie DA, Tailford LE, Gunning AP, Walshaw J, et al. Structural basis for adaptation of lactobacilli to gastrointestinal mucus. Environ Microbiol. 2014;16(3):888–903. doi: 10.1111/1462-2920.12377.

5. Nishiyama K, Sugiyama M, Mukai T. Adhesion properties of lactic acid bacteria on intestinal mucin. Microorganisms. 2016;4(3). doi: 10.3390/microorganisms4030034.

6. Savijoki K, Nyman TA, Kainulainen V, Miettinen I, Siljamäki P, Fallarero A, et al. Growth mode and carbon source Impact the surfaceome dynamics of Lactobacillus rhamnosus GG. Front Microbiol. 2019;10:1272. doi: 10.3389/fmicb.2019.01272.

7. Hynönen U, Palva A. Lactobacillus surface layer proteins: structure, function and applications. Appl Microbiol Biotechnol. 2013;97(12):5225–5243. doi: 10.1007/s00253-013-4962-2.

8. Konstantinov SR, Smidt H, de Vos WM, Bruijns SC, Singh SK, Valence F, et al. S layer protein A of Lactobacillus acidophilus NCFM regulates immature dendritic cell and T cell functions. Proc Natl Acad Sci U S A. 2008;105(49):19474–19479. doi: 10.1073/pnas.0810305105.

9. Yamasaki-Yashiki S, Sawada H, Kino-Oka M, Katakura Y. Analysis of gene expression profiles of Lactobacillus paracasei induced by direct contact with Saccharomyces cerevisiae through recognition of yeast mannan. Biosci Microbiota Food Health. 2017;36(1):17–25. doi: 10.12938/bmfh.BMFH-2016-015.

10. Hirayama S, Furukawa S, Ogihara H, Morinaga Y. Yeast mannan structure necessary for co-aggregation with Lactobacillus plantarum ML11-11. Biochem Biophys Res Commun. 2012;419(4):652–655. doi: 10.1016/j.bbrc.2012.02.068.

11. Nejati F, Junne S, Neubauer P. A big world in small grain: a review of natural milk kefir starters. Microorganisms. 2020;8(2). doi: 10.3390/microorganisms8020192.

12. Klijn N, Weerkamp AH, de Vos WM. Genetic marking of Lactococcus lactis shows its survival in the human gastrointestinal tract. Appl Environ Microbiol. 1995;61(7):2771–2774. doi: 10.1128/aem.61.7.2771-2774.1995.

13. Kimoto-Nira H. New lactic acid bacteria for skin health via oral intake of heat-killed or live cells. Anim Sci J. 2018;89(6):835–842. doi: 10.1111/asj.13017.

14. Sugimura T, Jounai K, Ohshio K, Tanaka T, Suwa M, Fujiwara D. Immunomodulatory effect of Lactococcus lactis JCM5805 on human plasmacytoid dendritic cells. Clin Immunol. 2013;149(3):509–518. doi: 10.1016/j.clim.2013.10.007.

15. Nomura M, Kobayashi M, Narita T, Kimoto-Nira H, Okamoto T. Phenotypic and molecular characterization of Lactococcus lactis from milk and plants. J Appl Microbiol. 2006;101(2):396–405. doi: 10.1111/j.1365-2672.2006.02949.x.

16. Fallico V, McAuliffe O, Fitzgerald GF, Ross RP. Plasmids of raw milk cheese isolate Lactococcus lactis subsp. lactis biovar diacetylactis DPC3901 suggest a plant-based origin for the strain. Appl Environ Microbiol. 2011;77(18):6451–6462. doi: 10.1128/AEM.00661-11.

17. Kimoto H, Mizumachi K, Okamoto T, Kurisaki J. New Lactococcus strain with immunomodulatory activity: enhancement of Th1-type immune response. Microbiol Immunol. 2004;48(2):75–82. doi: 10.1111/j.1348-0421.2004.tb03490.x.

18. Suzuki C, Kimoto-Nira H, Kobayashi M, Nomura M, Sasaki K, Mizumachi K. Immunomodulatory and cytotoxic effects of various Lactococcus strains on the murine macrophage cell line J774.1. Int J Food Microbiol. 2008;123(1-2):159–165. doi: 10.1016/j.ijfoodmicro.2007.12.022.

19. Nakano K, Minami M, Shinzato M, Shimoji M, Ashimine N, Shiroma A, et al. Complete genome sequence of Lactococcus lactis subsp. lactis G50 with immunostimulating activity, isolated from Napier grass. Genome Announc. 2018;6(8). doi: 10.1128/genomeA.00069-18.

20. Otto R, Tenbrink B, Veldkamp H, Konings WN. The relation between growth rate and electrochemical proton gradient of Streptococcus cremoris. FEMS Microbiology Letters. 1983;16(1):69–74.

21. Suzuki C, Kobayashi M, Kimoto-Nira H. Novel exopolysaccharides produced by Lactococcus lactis subsp. lactis, and the diversity of epsE genes in the exopolysaccharide biosynthesis gene clusters. Biosci Biotechnol Biochem. 2013;77(10):2013–2018. doi: 10.1271/bbb.130322.

22. Tanaka M, Odani T, Yuuki T, Ohtake Y, Shimma Y-I, Yoko-O T, et al. Yeast strain releasing mannan protein and method of producing mannan protein WO2006025295A1 2005.

23. Nomura M, Kobayashi M, Okamoto T. Rapid PCR-based method which can determine both phenotype and genotype of Lactococcus lactis subspecies. Appl Environ Microbiol. 2002;68(5):2209–2213. doi: 10.1128/AEM.68.5.2209-2213.2002.

24. Nomura M, Kimoto H, Someya Y, Suzuki I. Novel characteristic for distinguishing Lactococcus lactis subsp. lactis from subsp. cremoris. Int J Syst Bacteriol. 1999;49 Pt 1:163–166. doi: 10.1099/00207713-49-1-163.

25. Kimoto H, Nomura M, Suzuki I. Growth energetics of Lactococcus lactis subsp. lactis biovar diacetylactis in cometabolism of citrate and glucose. Int Dairy J. 1999;9(12):857–863.

26. Meyrand M, Guillot A, Goin M, Furlan S, Armalyte J, Kulakauskas S, et al. Surface proteome analysis of a natural isolate of Lactococcus lactis reveals the presence of pili able to bind human intestinal epithelial cells. Mol Cell Proteomics. 2013;12(12):3935–3947. doi: 10.1074/mcp.M113.029066.

27. Holm L, Rosenstrom P. Dali server: conservation mapping in 3D. Nucleic Acids Res. 2010;38(Web Server issue):W545–549. doi: 10.1093/nar/gkq366.

28. Kabsch W. XDS. Acta Crystallogr D Biol Crystallogr. 2010;66(Pt 2):125–132. doi: 10.1107/S0907444909047337.

29. Evans P. Scaling and assessment of data quality. Acta Crystallogr D Biol Crystallogr. 2006;62(Pt 1):72–82. doi: 10.1107/S0907444905036693.

30. Winn MD, Ballard CC, Cowtan KD, Dodson EJ, Emsley P, Evans PR, et al. Overview of the CCP4 suite and current developments. Acta Crystallogr D Biol Crystallogr. 2011;67(Pt 4):235–242. doi: 10.1107/S0907444910045749.

31. Vagin A, Teplyakov A. Molecular replacement with MOLREP. Acta Crystallogr D Biol Crystallogr. 2010;66(Pt 1):22–25. doi: 10.1107/S0907444909042589.

32. Yang YH, Jiang YL, Zhang J, Wang L, Bai XH, Zhang SJ, et al. Structural insights into SraP-mediated Staphylococcus aureus adhesion to host cells. PLoS Pathog. 2014;10(6):e1004169. doi: 10.1371/journal.ppat.1004169.

33. Emsley P, Cowtan K. Coot: model-building tools for molecular graphics. Acta Crystallogr D Biol Crystallogr. 2004;60(Pt 12 Pt 1):2126–2132. doi: 10.1107/S0907444904019158.

34. Murshudov GN, Skubak P, Lebedev AA, Pannu NS, Steiner RA, Nicholls RA, et al. REFMAC5 for the refinement of macromolecular crystal structures. Acta Crystallogr D Biol Crystallogr. 2011;67(Pt 4):355–367. doi: 10.1107/S0907444911001314.

35. Chen VB, Arendall WB, 3rd, Headd JJ, Keedy DA, Immormino RM, Kapral GJ, et al. MolProbity: all-atom structure validation for macromolecular crystallography. Acta Crystallogr D Biol Crystallogr. 2010;66(Pt 1):12–21. doi: 10.1107/S0907444909042073.

36. Biswas I, Gruss A, Ehrlich SD, Maguin E. High-efficiency gene inactivation and replacement system for Gram-positive bacteria. J Bacteriol. 1993;175(11):3628–3635. doi: 10.1128/jb.175.11.3628-3635.1993.

37. Oxaran V, Ledue-Clier F, Dieye Y, Herry JM, Pechoux C, Meylheuc T, et al. Pilus biogenesis in Lactococcus lactis: molecular characterization and role in aggregation and biofilm formation. PLos One. 2012;7(12):e50989. doi: 10.1371/journal.pone.0050989.

38. Maguin E, Prevost H, Ehrlich SD, Gruss A. Efficient insertional mutagenesis in lactococci and other Gram-positive bacteria. J Bacteriol. 1996;178(3):931–935. doi: 10.1128/jb.178.3.931-935.1996.

39. Papagianni M, Avramidis N, Filioussis G. High efficiency electrotransformation of Lactococcus lactis spp. lactis cells pretreated with lithium acetate and dithiothreitol. BMC Biotechnol. 2007;7:15. doi: 10.1186/1472-6750-7-15.

40. Dieye Y, Oxaran V, Ledue-Clier F, Alkhalaf W, Buist G, Juillard V, et al. Functionality of sortase A in Lactococcus lactis. Appl Environ Microbiol. 2010;76(21):7332–7337. doi: 10.1128/AEM.00928-10.

41. Kinoshita H, Ohuchi S, Arakawa K, Watanabe M, Kitazawa H, Saito T. Isolation of lactic acid bacteria bound to the porcine intestinal mucosa and an analysis of their moonlighting adhesins. Biosci Microbiota Food Health. 2016;35(4):185–196. doi: 10.12938/bmfh.16-012.

42. Amblee V, Jeffery CJ. Physical features of intracellular proteins that moonlight on the cell surface. PLoS One. 2015;10(6):e0130575. doi: 10.1371/journal.pone.0130575.

43. MacKenzie DA, Tailford LE, Hemmings AM, Juge N. Crystal structure of a mucus-binding protein repeat reveals an unexpected functional immunoglobulin binding activity. J Biol Chem. 2009;284(47):32444–32453. doi: 10.1074/jbc.M109.040907.

44. Arras SDM, Hibbard TR, Mitsugi-McHattie L, Woods MA, Johnson CE, Munkacsi A, et al. Creeping yeast: a simple, cheap and robust protocol for the identification of mating type in Saccharomyces cerevisiae. FEMS Yeast Res. 2022;22(1). doi: 10.1093/femsyr/foac017.

45. Suzuki C, Shimma YI. P-type ATPase spf1 mutants show a novel resistance mechanism for the killer toxin SMKT. Mol Microbiol. 1999;32(4):813–823. doi: 10.1046/j.1365-2958.1999.01400.x.

46. Pretzer G, Snel J, Molenaar D, Wiersma A, Bron PA, Lambert J, et al. Biodiversity-based identification and functional characterization of the mannose-specific adhesin of Lactobacillus plantarum. J Bacteriol. 2005;187(17):6128–6136. doi: 10.1128/JB.187.17.6128-6136.2005.

47. von Ossowski I, Satokari R, Reunanen J, Lebeer S, De Keersmaecker SC, Vanderleyden J, et al. Functional characterization of a mucus-specific LPXTG surface adhesin from probiotic Lactobacillus rhamnosus GG. Appl Environ Microbiol. 2011;77(13):4465–4472. doi: 10.1128/AEM.02497-10.

48. Camacho C, Coulouris G, Avagyan V, Ma N, Papadopoulos J, Bealer K, et al. BLAST+: architecture and applications. BMC Bioinformatics. 2009;10:421. doi: 10.1186/1471-2105-10-421.

49. Berlec A, Zadravec P, Jevnikar Z, Štrukelj B. Identification of candidate carrier proteins for surface display on Lactococcus lactis by theoretical and experimental analyses of the surface proteome. Appl Environ Microbiol. 2011;77(4):1292–1300. doi: 10.1128/AEM.02102-10.

50. Yanagida F, Chen YS, Shinohara T. Searching for bacteriocin-producing lactic acid bacteria in soil. J Gen Appl Microbiol. 2006;52(1):21–28. doi: 10.2323/jgam.52.21.

51. Siezen RJ, Starrenburg MJ, Boekhorst J, Renckens B, Molenaar D, van Hylckama Vlieg JE. Genome-scale genotype-phenotype matching of two Lactococcus lactis isolates from plants identifies mechanisms of adaptation to the plant niche. Appl Environ Microbiol. 2008;74(2):424–436. doi: 10.1128/AEM.01850-07.

