## Supplementary material for "Cell-surface protein YwfG of *Lactococcus lactis* binds to α-1,2-linked mannose": S1 Table

S1 Table Plasmids and primers used in this study.

| Plasmid | Relevant characteristics | Reference |
| --- | --- | --- |
| pG+host9 | Ts derivative of pWV01, Erythromycin^r^ | [38] |
| pGEM-T Easy | ColE1, Ampicillin (Amp)^r^ | Promega |
| pCS336 | 963 bp upstream of *ywfG* cloned into pGEM-T Easy, Amp^r^ | This study |
| pCS337 | 990 bp downstream of *ywfG* cloned into pGEM-T Easy, Amp^r^ | This study |
| pCS338 | 963 bp upstream of *ywfG* and 990 bp downstream of *ywfG* cloned into pGEM-T Easy, Amp^r^ | This study |
| pCS339 | pCS338 and pG+host9 were ligated at SacII site | This study |
| Primer | Sequence | |
| Fw28 | 5′-CCGCGCGGCAGCCATATGCCCGACTCTTTTAAAATACAACGG-3 | |
| Rv270 | 5′-GAGCTCGAATTCGGATCCTCAACCTTGAGCTACCGTATAAGT-3′ | |
| Rv336 | 5′-GAGCTCGAATTCGGATCCTCAATTTCGAGTATAAACATAGTT-3′ | |
| Rv511 | 5′-GAGCTCGAATTCGGATCCTCAGTTACGTTTATAAACGACCTT-3′ | |
| Fw860 | 5′-CCGCGCGGCAGCCATATGCAAGGAACCATTGATGTTACTTAT-3′ | |
| Rv1034 | 5′-GAGCTCGAATTCGGATCCTCAAGGCACCTTATGATAAACGAC-3′ | |
| 5’G50_5140 | 5′-TACTACAAAAAGAGAGGAAAAGGGG-3′ | |
| 3’G50_6123 | 5′-CTGTGTTACTTTTTGCTTTCGCTCG-3′ | |
| 5’G50_1026 | 5′-TTTCTTGGGTTTTGGGATCTTGAAC-3′ | |
| 3’G50_3016 | 5′-GTCGTTTATCATAAGGTGCCTGCAG-3′ | |
