## Supporting Figs for "Cell-surface protein YwfG of *Lactococcus lactis* binds to α-1,2-linked mannose"

A

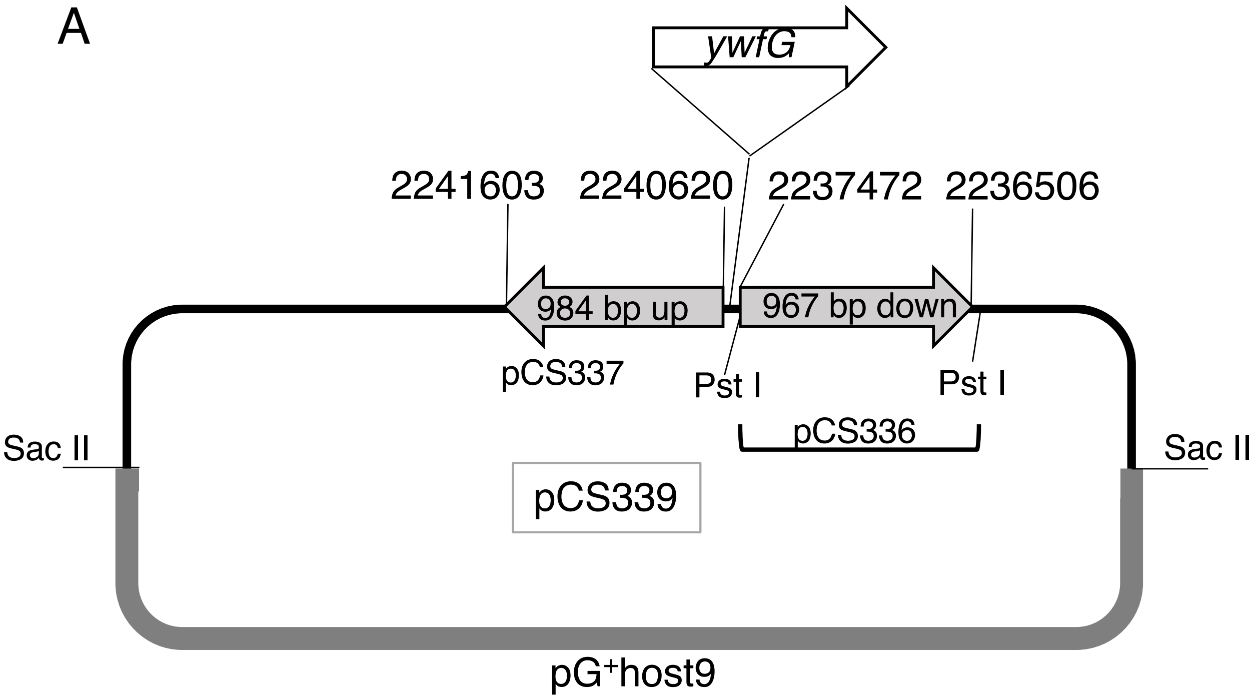
ywfG
2241603
2240620
2237472
2236506
984 bp up
967 bp down
pCS337
Pst I
Pst I
pCS336
Sac II
Sac II
pCS339
pG+host9

B

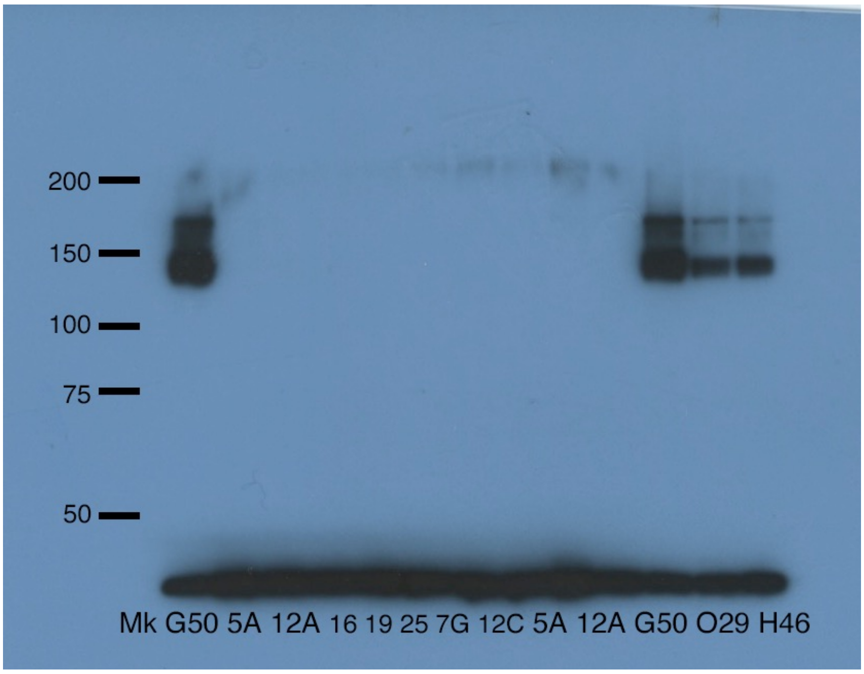
200
150
100
75
50
Mk G50 5A 12A 16 19 25 7G 12C 5A 12A G50 O29 H46

S1 Fig

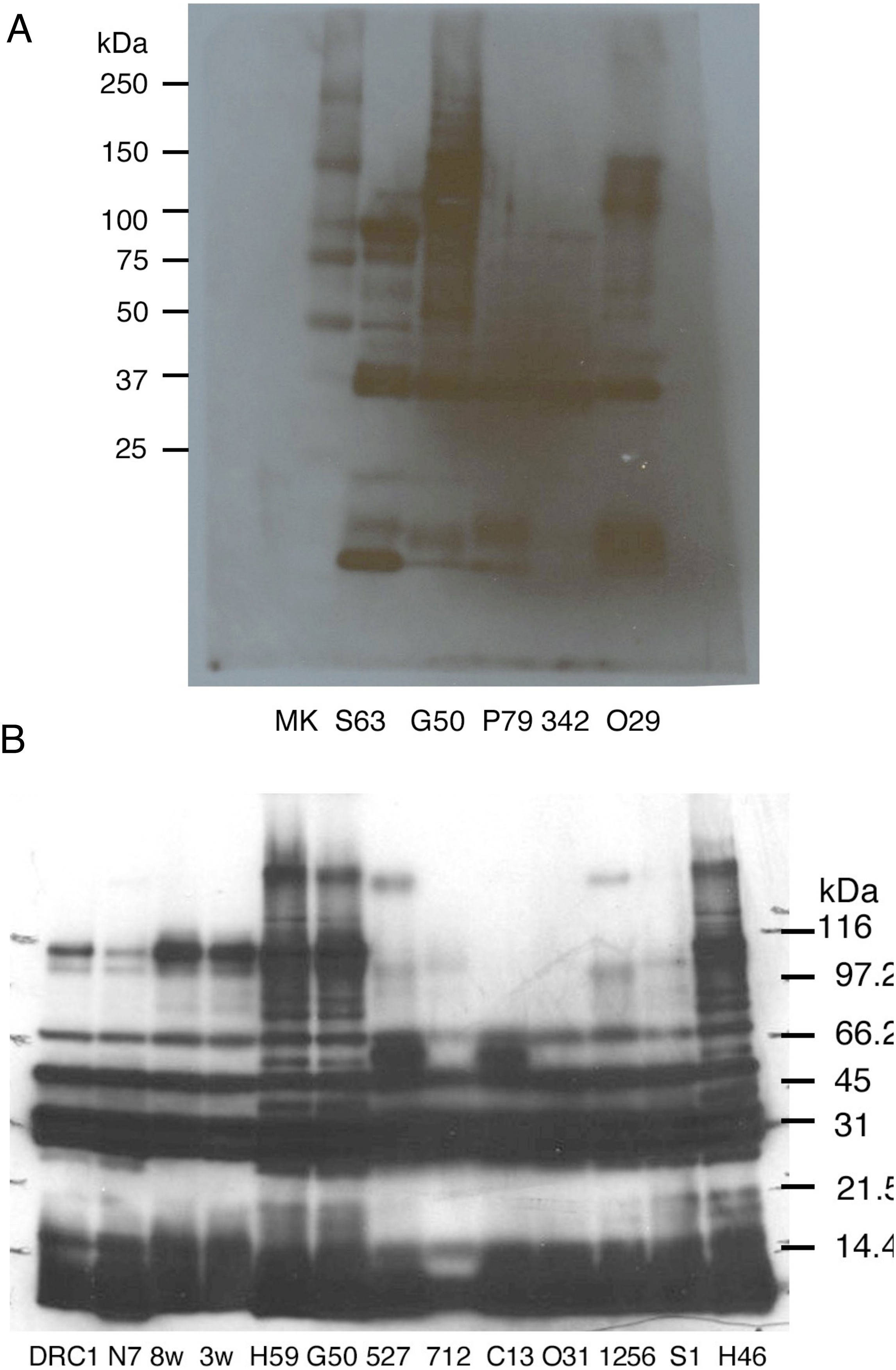
A
kDa
250
150
100
75
50
37
25
MK S63 G50 P79 342 O29
B
kDa
116
97.2
66.2
45
31
21.5
14.4
DRC1 N7 8w 3w H59 G50 527 712 C13 O31 1256 S1 H46

S2 Fig

MNKKSSALLTM**GTLTLLAGGGVVLTNLPDSFKIQR**VYAATSRDITVYPK**DFLTYFQRNGS** 60

**AAGFDYDLATYTQTLTPNKASQAGNVTLKTKVDMSQNFTFTGK**INLGDKAQNAGGADGVG 120

FLFHPGDTNVVGAPGGAAGIGGVNGAFGFKLDTYYNGVGENSFTPDPSNFK**GKPFGAFVD** 180

**GLNGQAK**TIASSAQSISEPSNNNFVDFTMSYNGATKVMSVTYGGQTWTQDVSSFVGTNQA 240

MSFSIAASTGAFMNLQQLRNVNFTYTVAQGTVIANYVDEQGNTIAQQETTSGDIDTPYVT 300

SQK**TIPGYTFKASNGAATSGNYAANDQTVNYVYTRNQGSIDVTYIDQTTGQTLSKK**DLSG 360

GTGDSSNYTTADTIKSYTDAGYELVSDNYPSGGTVFTDTAQHYVVNLK**QKLVVSSEQKQV** 420

**NETIQYVVYEDGSKAADDYNAPPLNFTRSVTTNQVTGEK**TYGDWQAQNGDSFGEVVSPTIK 480

GETADQLK**IDAISGITANSADIQK**KVVYKR**NQGTIDVTYIDETTGQVLTKKDLSGGTDDP** 540

**SNYTTADDIK**SYTAK**GYELVSDDYPSGGTVFTDEPQHYVVK**LK**HGLTESTDKKAVNQVIH** 600

**YVYEGGGEAATDHNATVDFSR**TITTDRVTNDKTYGDWTADNGDSFASVTSPVIDGYTADQ 660

LK**VSEMTGITADTEDISVTVTYTR**NQGTIDITYIDQTTGQTLSEKDLSGGTGDDSGYTTA 720

DTIK**SYTDKGYELVSNDYPEDGTKFADDPQHYIVRLKHGLTEVTENK**TVNQVIHYVYEGG 780

GEAATDHNATVVFSQTITTDKVTGEKTYSDWTADNGDSFASVTSPVIDGYTADQLKVSEM 840

TGITVDTEDISVTVTYTR**NQGTIDVTYIDETTGK**ILTTK**DLSGGTGDDSGYTTADTIKSY** 900

**TDKGYELVSNDYPEDGTKFADDPQHYIVRLKHGTVVETENKSVNEVIHYVYDNGDKAADN** 960

**YKATIVFSRTITTDKVTGEKTYSDWTADNGGR**FAAVLSPIIKDYIASQLK**IDEMTGITVD** 1020

**TADIERIVVYHKVPAGIIVPPVHPDKPSQPSNNQSK**TPTAKAVK**DSKPTDVLPSTGDSQK** 1080

SQIVLTLLGIMAVIISPLALLLRRRKQ 1107

S3 Fig

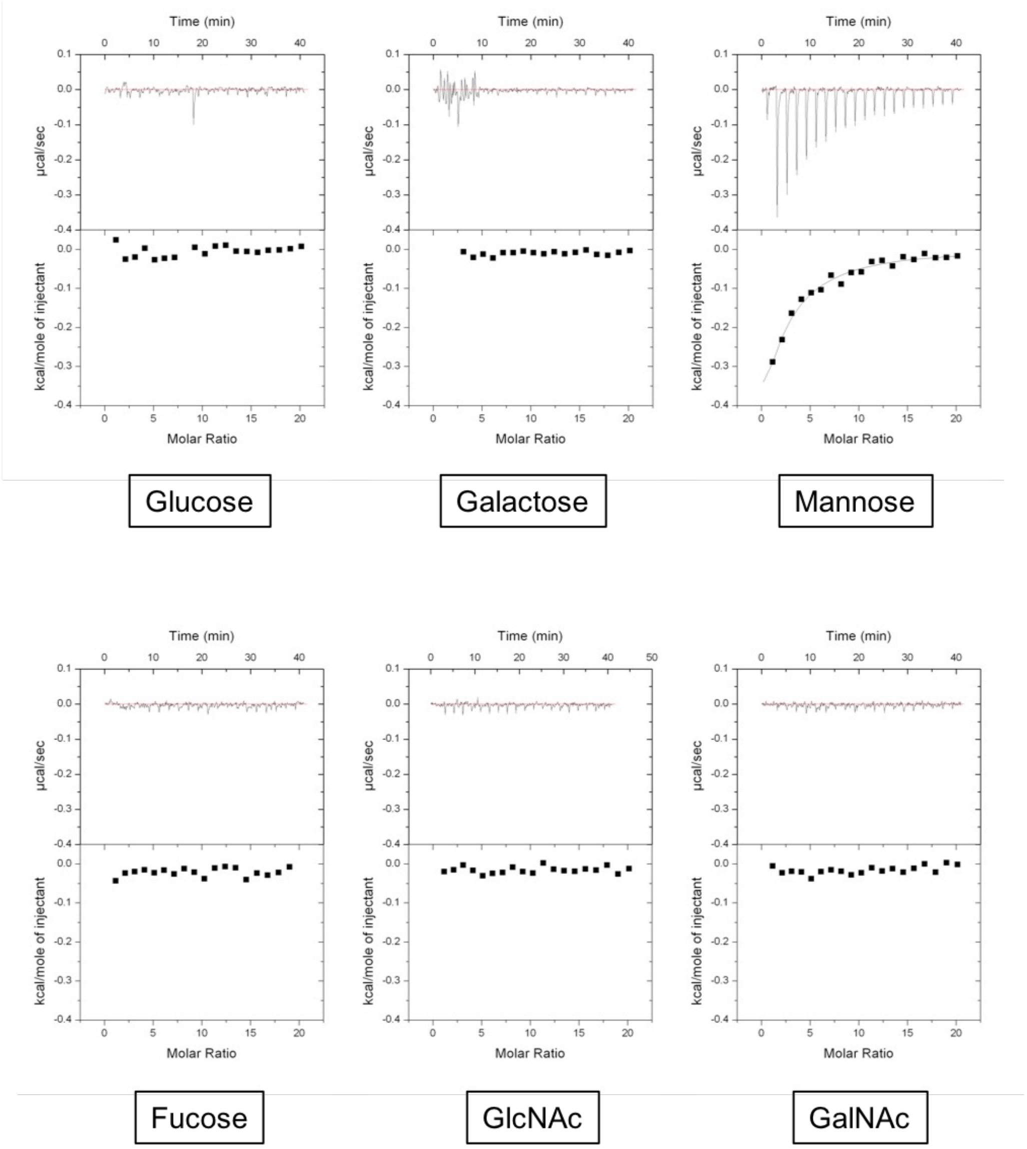
Time (min)
µcal/sec
kcal/mole of injectant
Molar Ratio
Glucose
Galactose
Mannose
Fucose
GlcNAc
GalNAc

S4 Fig

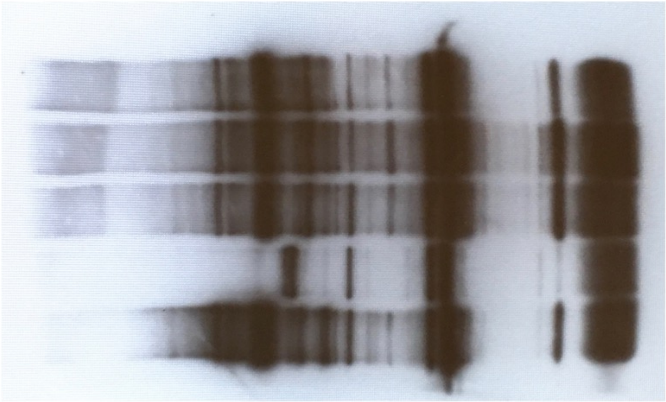

Western blotting (Figure 1A)

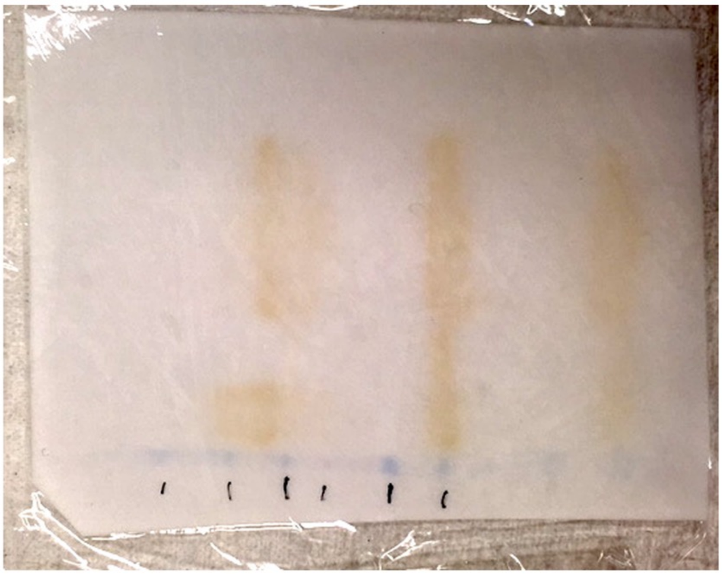

Blot membrane after exposure

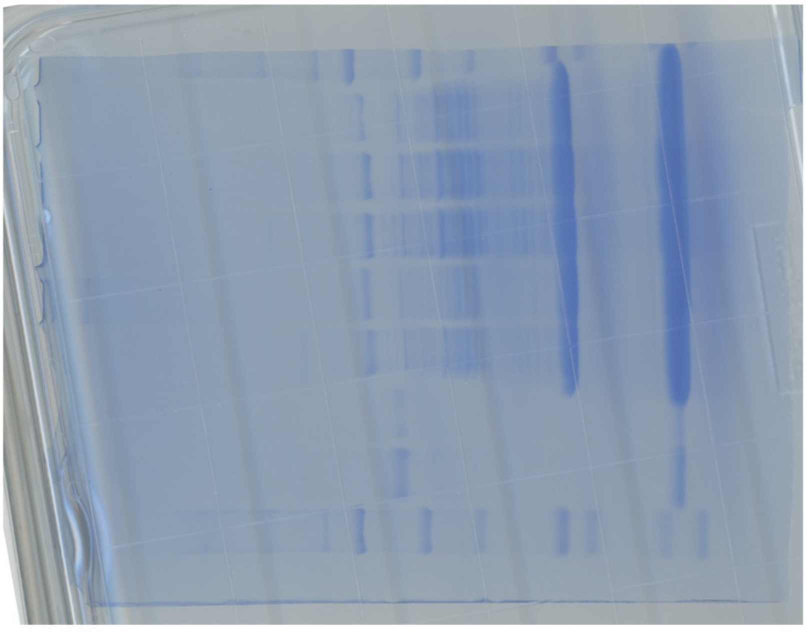

CBB staining (Figure 1B)

S1_raw_Fig 1 image

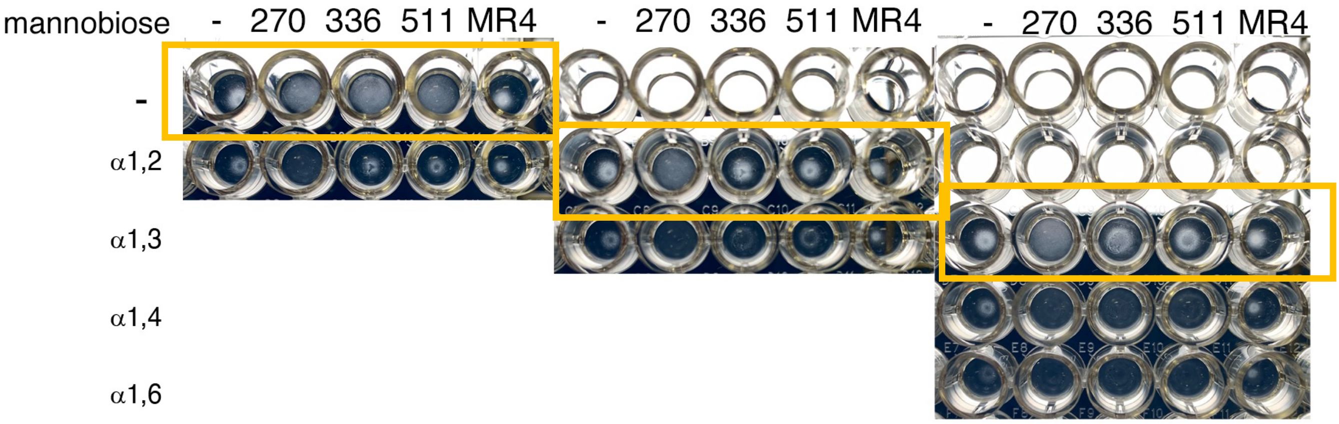


S2_raw_Fig 7C image
